# RIPOR2 promotes multinucleation of melanoma cells downstream of the RAS/ERK oncogenic pathway

**DOI:** 10.1101/2025.06.26.661696

**Authors:** Axelle Wilmerding, Aurélie Richard, Nicolas Macagno, Estelle Hirsinger, Tarek Gharsalli, Léa Bellenger, Naïra Naouar, Caroline Gaudy, Stéphanie Mallet, Nathalie Degardin, Lauranne Bouteille, Nathalie Caruso, Delphine Duprez, Yacine Graba, Souhila Medjkane, Heather C. Etchevers, Marie-Claire Delfini

## Abstract

One-third of skin melanomas arise from melanocytic nevi, benign skin lesions composed of clustered melanocytes. Benign nevi are associated with overactivation of the mitogen-activated protein kinase RAS/ERK pathway, resulting from driver mutations, most commonly in the *BRAF* or *NRAS* gene. However, this overactivation in melanocytes is insufficient to induce melanoma formation, as only a minority of benign nevi give rise to melanoma. Overactivation of the RAS/ERK pathway promotes genetic and epigenetic alterations by inducing aneuploidy, but the processes by which nevi evolve into melanoma via RAS/ERK pathway-dependent aneuploidy are only partially understood. Using single-nucleus RNA sequencing after overactivation of the RAS/ERK pathway in the chicken embryo, we discovered that *RIPOR2* is a positive transcriptional target of this pathway, including in melanocyte precursors. Similar transcriptional control of *RIPOR2* by RAS/ERK is conserved in human melanoma cells. *RIPOR2* emerged as an attractive target because it encodes an atypical RHOA inhibitory protein involved in the development of physiologically multinucleated cell types. Multinucleation in cancer has been shown to promote aneuploidy, which correlates with tumor aggressiveness. We found that RIPOR2 is ectopically expressed in human nevi and skin melanomas and functionally promotes multinucleation in both an animal model and human tumor-derived cells, including melanoma cell lines. Our results suggest that RIPOR2 expression, downstream of RAS/ERK overactivation in skin melanocytes, promotes the emergence of multinucleated cells, a previously overlooked step in melanoma formation.

## Introduction

Cutaneous melanoma is aggressive: although it only accounts for 5% of all skin cancers, it is the leading cause of skin cancer-related deaths. Approximately one-third of melanomas arise from benign melanocytic nevi, neoplasms of melanocytes, the skin’s normal pigment-producing cells^1^. During melanoma development, the RAS/RAF/MEK/ERK (RAS/ERK) pathway in melanocytes is overactivated in more than 80% of cutaneous melanomas ^2^. Genomic and transcriptomic analyses have revealed that RAS/ERK signaling is overactivated in nevi and melanomas from the earliest stages of neoplasia, with melanocyte clones reflecting individual driver mutations ^3^. From the early neoplasm stage, overactivation of the RAS/ERK pathway correlates with tumor progression as malignant transformation advances ^4–6^. Although overactivation of the RAS/ERK pathway is central to melanoma development, it is not directly sufficient to induce malignancy since most nevi remain benign ^7^. The process of nevus progression to melanoma is only partially understood.

Many molecular effectors of the RAS/ERK pathway have been identified for decades, but its modulators continue to be discovered. Ligand stimulation of cell surface receptors, usually of the tyrosine kinase type, induces the conversion of RAS homologs (such as HRAS, NRAS, KRAS) into active forms, leading to the recruitment of homodimers or heterodimers of the RAF family (ARAF, BRAF, or CRAF, also known as RAF1) to the plasma membrane. RAF kinases phosphorylate MEKs (MEK1 and MEK2), which in turn phosphorylate two ERKs (ERK1 and ERK2) to transduce signals to hundreds of cytosolic and nuclear substrates ^8–10^. While the effects of oncogenic RAS signaling on cell cycle G1/S checkpoint clearance are well known, there is also evidence that RAS/ERK activation promotes aneuploidy ^11–16^. Several lines of evidence now support the argument that aneuploidy promotes tumorigenesis rather than being a transient event ^13–19^. In particular, enforced nuclear localization of MEK1, which leads to overactivation of ERK1/2 signaling, is sufficient to induce polyploidization and neoplastic transformation of cells *in vivo* ^14^, demonstrating that early oncogenic RAS/ERK activation contributes to tumor progression to melanoma via inducing aneuploidy.

The mechanisms by which oncogenic activation of the RAS/ERK pathway contributes to aneuploidy and melanoma progression remain poorly understood. Defects in mitosis/cytokinesis and non-physiological cell-cell fusion, resulting in multinucleated cells, have been shown to give rise to aneuploidy ^20,21^. Interestingly, in cutaneous melanoma, constitutive overactivation of the RAS/ERK pathway promotes the formation of senescent-like multinucleated cells by promoting cell-cell fusion and fragmentation following mitotic defects that confer better cell survival ^15^. *In vitro*, long-term expression of NRAS^Q61K^ in melanocytes also triggers a strong senescent-like phenotype associated with multinucleation, followed by the generation of tumor-initiating cells with stem cell-like properties ^16^.

In a RAS/ERK-induced neoplasm model, we generated multinucleated cells *in vivo* ^22^. This model consists of expression of a constitutively active form of MEK1 (MEK1ca) in the developing spinal cord of two-day-old chicken embryos. By targeting this tissue and stage, dorsal midline neural crest cells were transfected, including progenitors of all cutaneous melanocytes from both the dorsolateral migratory pathway and the later emerging multipotent Schwann cell precursors. Multinucleated cells appeared *de novo* in this neoplasia model as early as two days after electroporation, showing that induction of multinucleation is an early consequence of RAS/ERK overactivation in the multipotent neuroectoderm.

In this study, using single-nucleus RNA sequencing (snRNAseq), we discovered that the *RIPOR2* gene (RHO family-interacting cell polarization regulator 2, previously known as *FAM65B*) is a transcriptional target of the oncogenic RAS/ERK pathway. Furthermore, RIPOR2 is ectopically expressed in congenital human melanocytic nevi and melanomas. Its gain-of-function promotes multinucleation *in vivo* in chicken embryos and in human cell cultures, including in melanoma cells. These correlations suggest that RIPOR2 expression in nevi, triggered by overactivation of RAS/ERK by driver mutations, may contribute to the transition from nevus to melanoma by causing the appearance of abnormally multinucleated cells.

## RESULTS

### MEK1ca induces a transcriptional program overriding positional identity in the chicken neural tube

To identify RAS/ERK pathway effectors that may control the oncogenic progression of melanoma, we used an *in vivo* tumor model induced by MEK1ca-mediated ERK activation in the neural tube of two-day-old chicken embryos ^22^. At this stage, the trunk neural tube is composed of neural progenitors of the future spinal cord and premigratory neural crest cells in its most dorsal part, which will give rise to all cutaneous melanocytes ^23^. To isolate the transcriptional response of neural crest cells and their immediate progeny to MEK1ca expression, we performed single-nucleus RNA sequencing (snRNAseq) to compare the effects of electroporating a vector co-expressing MEK1ca and GFP with those of a vector expressing only GFP. One day after bilateral electroporation of the trunk neural tube, the entire body segment was dissected at the level of electroporation (Fig. 1A). Twenty-four embryos were pooled for each condition, yielding 17,070 nuclei for analysis by snRNAseq, of which 7,098 nuclei (168 GFP+ and 6,930 GFP-) were from the MEK1ca condition and 9,972 nuclei (394 GFP+ and 9,578 GFP-) from the control condition. The two datasets were merged using Seurat (Fig. 1B). We analyzed the dataset at a resolution of 0.6, resulting in 23 clusters.

**Figure 1:**
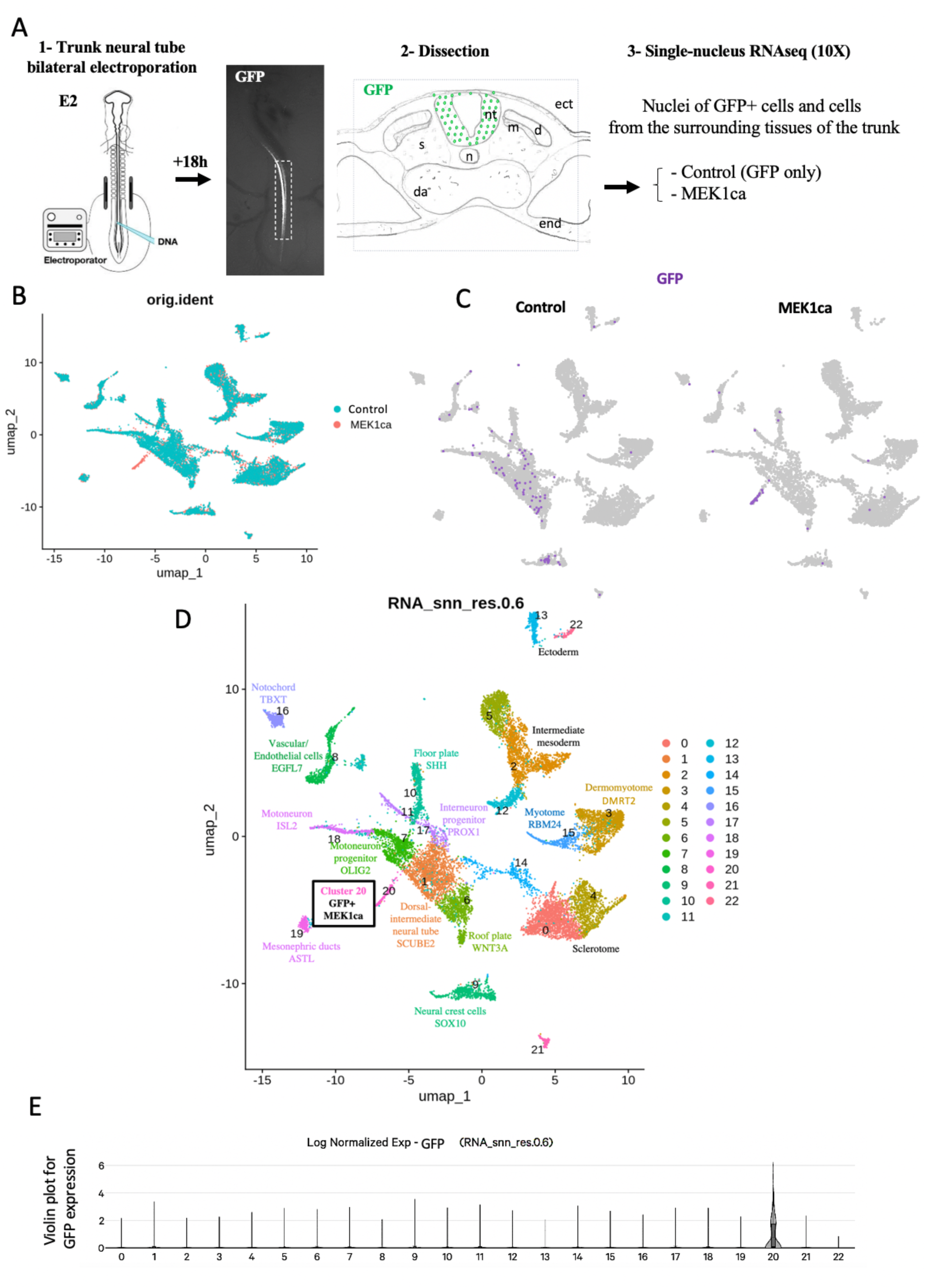
Neural tube cells expressing MEK1ca acquire a common, new molecular signature independent of their initial position. **A-** One day after bilateral electroporation of the neural tube of two-day old chicken embryos with PCIG (control vector expressing only GFP) or PCIG-MEK1ca (co-expressing the constitutively active form of MEK1 in addition to GFP), embryos were dissected at the electroporated level. snRNAseq analysis was conducted on nuclei of transfected (GFP+) cells as well as on nuclei from the surrounding tissues of the trunk (excluding therefore the head and the tail bud) (nt, neural tube; n, notochord; ect, ectoderm; da, dorsal aorta; m, myotome; dm, dermomyotome; s, sclerotome; end, endoderm). **B-** UMAP plot of the merged dataset (17,070 nuclei in total, including 9972 cells (394 GFP+ and 9578 GFP-) from the control condition, and 7098 cells (168 GFP+ and 6930 GFP-) from the MEK1ca condition.), showing distribution of each datasets of origin. **C-** Feature plots showing the distribution of GFP positive cells (GFP>1) in the control and MEK1ca datasets. **D-** An unsupervised UMAP subdivides cells within the trunk into 23 clusters at snn_res.0.6). **E-** Violin plots for GFP expression showing the distribution of transfected cells in each cluster.

A small population of 201 cells, corresponding to cluster 20, was found only in the MEK1ca sample (Fig. 1B-D). Feature plots of GFP expression (the transfection marker) showed that this cluster consisted of MEK1ca-transfected cells (Fig. 1C). All clusters were annotated using differentially expressed markers of major cell types in the three-day-old chicken embryo trunk (Fig. 1D and Supplementary Fig. 1). Among these, we annotated six neural tube clusters: cluster 1 (dorsal-intermediate neural tube, preferentially expressing *SCUBE2* ^24^), cluster 6 (*WNT3A+* roof plate ^25^, containing neural crest/melanocyte progenitors), cluster 7 (motoneuron progenitors, *OLIG2+* ^26^), cluster 10 (floor plate, *SHH+* ^27^), cluster 17 (interneuron progenitors, *PROX1+* ^28^), and cluster 18 (definitive motoneurons, *ISL2+* ^29^) (Fig. 1D and Supplementary Fig. 1B-G). Migrating neural crest cells of the trunk constituted cluster 9 (differentially expressing *SOX10* ^30^, Supplementary Fig. 1H). Among additional cell types, we identified the notochord (cluster 16, *TBXT+* ^31^, Supplementary Fig. 1I) , vascular/endothelial cells (cluster 8, *EGFL7+* ^32^, Supplementary Fig. 1J), mesonephric duct cells (*ASTL+* ^33^, Supplementary Fig. 1K), intermediate mesoderm (clusters 2, 5, and 12, *WT1+* ^34^, Supplementary Fig. 1L), sclerotome (clusters 0 and 4, *PAX1+* ^35^, Supplementary Fig. 1M), ectoderm (clusters 13 and 22, WNT6+ ^36^, Supplementary Fig. 1N), myotome (cluster 15, *RBM24*+ ^37^, Supplementary Fig. 1O), and dermomyotome (cluster 3, DRMT2 ^38^, Supplementary Fig. 1P) (Fig. 1D and Supplementary Fig. 1).

In the control condition, GFP-positive cells one day after electroporation were assigned in the UMAP to all six neural tube clusters as well as to cluster 9, corresponding to migrating neural crest cells (Fig. 1C-D). In contrast, while GFP-positive cells of the MEK1ca condition in the embryo were physically present in the neural tube ^22^, their snRNAseq assignment to the new cluster 20 (Fig. 1C-E) indicates that neural tube cells (including neural crest-derived melanocyte progenitors) respond similarly to MEK1ca. Furthermore, their collective transcriptome indicates a novel cell type.

### Analysis of differential gene expression after MEK1ca expression in the chicken embryo neural tube using snRNAseq is highly consistent with bulk RNAseq data

Differentially expressed genes (DEGs) were assessed among GFP-expressing cells (GFP>1) in the MEK1ca and control conditions. 424 upregulated and 528 downregulated transcripts were distinguished (Supplementary Table 1). Of the 50 most upregulated genes (Fig. 2A), 86% (43/50) were also present among the DEGs when assessed in independent bulk RNAseq experiments (p-value <0.05) performed in the same model at the same stage ^22^. For example, *IL1R1*, the most significantly upregulated gene in bulk RNAseq ^22^, was also among the top three most significantly upregulated genes in this comparison (Fig. 2A). This suggests that MEK1ca cell-autonomously activates *IL1R1* in transfected cells, as confirmed by *in situ* hybridization after neural tube electroporation (Supplementary Fig. 2). *CDX4*, *IL17RD*, *CHST15,* and *TMEM132C* (Fig. 2A) were previously validated to be upregulated in the neural tube after MEK1ca electroporation by *in situ* hybridization ^22,39^. In conclusion, the snRNAseq experiment replicated and provided finer resolution than our previous bulk RNAseq analysis ^22^. The list of MEK1ca versus control DGEs among GFP-positive cells (Supplementary Table 1) represents potential RAS/ERK pathway effectors responsible for the multinucleation observed in the chicken embryo neural tube after MEK1ca expression.

**Figure 2:**
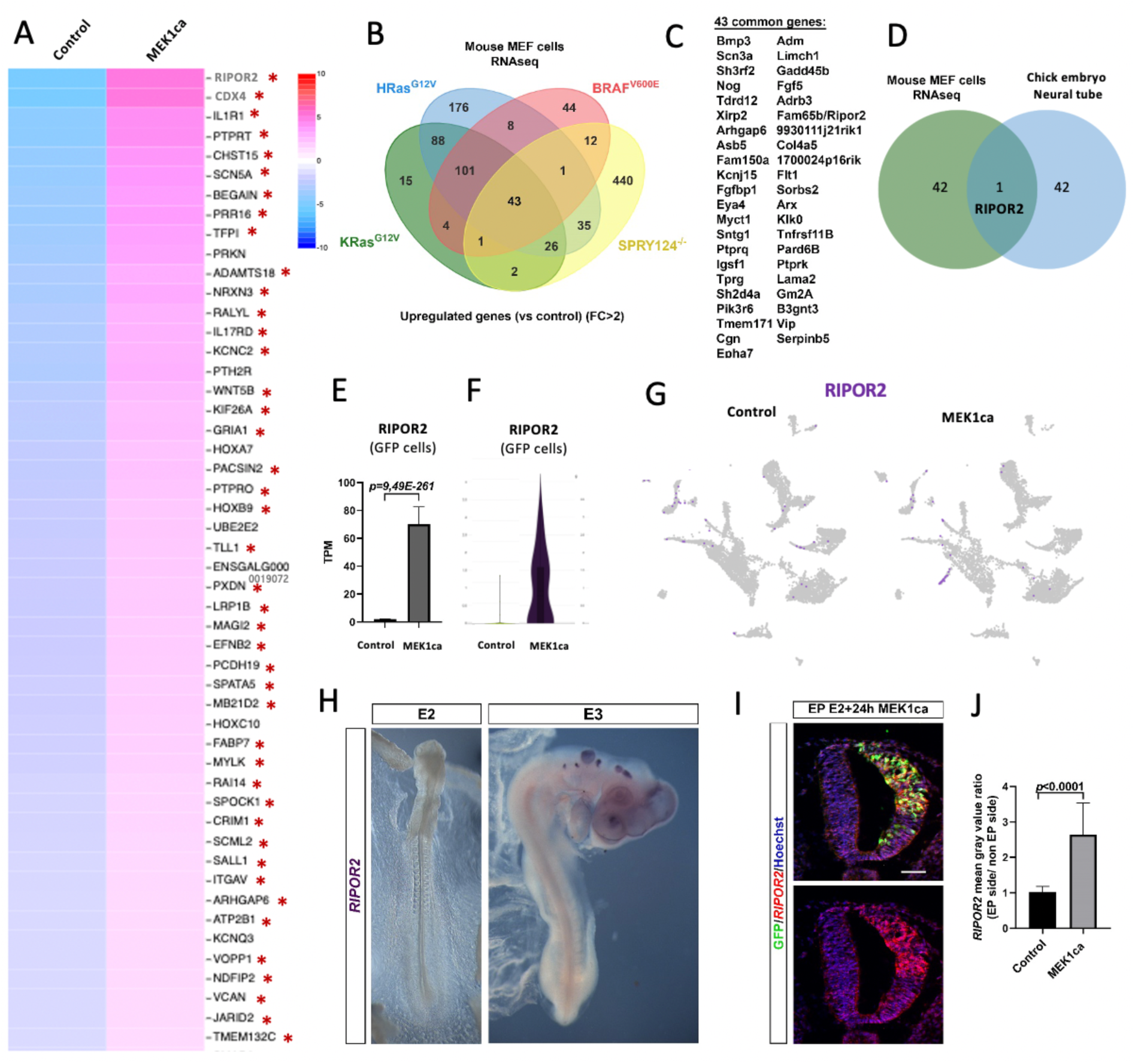
RIPOR2 is commonly upregulated across conditions of ERK overactivation in mouse and chicken embryos. **A –** Heat map from snRNAseq analysis showing the 50 most upregulated genes in transfected cells after MEK1ca expression (MEK1ca versus Control in GFP>1 cells). Genes with red stars (43/50) are in the list of upregulated genes in bulk RNAseq after MEK1ca expression (Wilmerding et al., 2022). **B -** Venn diagram of the upregulated genes (fold change>2) between 4 conditions (HRas^G12V^, KRas^G12V^, BRaf^V600E^ and Spry124−/−) in which RAS/ERK signaling is overactivated in MEFs (from Nabet et al., 2015). 43 genes **(C)** are commonly regulated between those 4 conditions. **D-** Venn diagram comparing the list of 43 upregulated genes in MAPK gain-of-function mouse models with the 43 genes upregulated in the chicken embryo neural tube after MEK1ca expression **(A)**. Only RIPOR2 is commonly deregulated. **E-** Mean expression of *RIPOR2* in TPM (transcripts per kilobase million), obtained for the two replicates of the control (pCIG) and MEK1ca in the chicken embryo (one day after electroporation) from bulk data (Wilmerding et al., 2022). **F-** Violin plot for *RIPOR2* expression in transfected/GFP expressing cells (GFP>1) for the two conditions from snRNAseq. **G-** Feature plots for *RIPOR2* expression in the control and MEK1ca datasets. **H-** Dorsal view of two- and three-day-old chicken embryo after a whole-mount *in situ* hybridization with the chicken probe *RIPOR2* **I-J-** Fluorescent *in situ* hybridization with a chicken *RIPOR2* probe and immunofluorescence with anti-GFP antibody on trunk transverse section of chicken embryo one day after electroporation of the MEK1ca plasmid, confirm the upregulation of *RIPOR2* by MEK1ca, which is statistically significant in the quantification compared to control (pCIG). (n=3 animals/18 sections, two-tailed Mann–Whitney test, error bars represent s.d.). Blue is Hoechst staining. Scale bar: 50µm.

### RIPOR2 is a general and conserved transcriptional target of the RAS/ERK oncogenic signaling pathway

Based on the gene list from the intersection of bulk and snRNAseq transcriptomic results (Fig. 2A), we identified RIPOR2 as a general and conserved RAS/ERK oncogenic transcriptional target. Indeed, we took advantage of published data that present transcriptomic results for four different conditions of RAS/ERK signal overactivation in mouse embryonic fibroblast MEF cells ^40^. Three of these murine conditions result from expression of mutant oncogenes (HRas^G12V^, KRas^G12V^, or BRaf^V600E^ expressing MEF cells), and the last is a knockout of Sprouty genes (Spry 1,2,4 ^-/-^ MEF cells) ^40^. Sprouty genes are part of the negative feedback loops that limit the activation of the RAS/RAF/MEK/ERK pathway. There is a high degree of overlap in genes deregulated in response to KRas^G12V^, HRas^G12V^, and BRaf^V600E^, but the genes modulated in Spry1,2,4^-/-^ are largely different ^40^. Applying a Fold Change > 2 filter to MEF transcriptomic data, only 43 of the 573 genes modulated in Spry1,2,4^-/-^ are also deregulated in response to KRas^G12V^, HRas^G12V^, and BRaf^V600E^ (7.5%) (Fig. 2B-C). Comparing these 43 genes commonly upregulated in these four conditions in mouse MEF cells (Fig. 2C) with the list of 43 upregulated genes after MEK1ca expression in chicken embryo neural tube (common between bulk and snRNAseq data, Fig. 2A) allowed us to identify RIPOR2 at the intersection (Fig. 2D). The particularly stringent conditions for identifying this gene resulted in identifying only one at the intersection of the five conditions in which the RAS/ERK pathway is overactivated, suggesting that the regulation of RIPOR2 by the RAS/ERK pathway is a conserved feature of RAS/ERK overactivation.

RIPOR2, or FAM65B (also known as PL48, MYONAP, and C6orf32) encodes for a protein that is upregulated during placenta and skeletal muscle differentiation, two multinucleated tissues involving cell fusion ^41,42^. It has been shown to induce the formation of membrane protrusions in muscle cells ^42^ and to act as an unusual RHOA regulator in T-lymphocytes ^43^ and neutrophils ^44^. In T-lymphocytes, RIPOR2 acts as a quiescence factor and can inhibit cell proliferation via mitotic spindle defects leading to mitotic failures, both in a T-cell line and transformed cell lines (such as HeLa cells) ^22, 43^. So far, no link between RIPOR2 and the RAS/ERK pathway has been described. The function of the RIPOR2 protein in solid tumors has not been described. Interestingly, it has been shown to be highly expressed in prostate cancer cells with stem cell-like properties ^45^, but its role in this cancer remains unknown.

The RIPOR2 gene is one of the most upregulated genes after MEK1ca expression in the chicken embryo neural tube, with a fold change >26 and padj= 3.9 x 10^-257^ in bulk data ^22^ (Fig. 2E). It is also at the top of the list on the heatmap of genes upregulated by MEK1ca according to snRNAseq data (Fig. 2A). The violin plot (in GFP-expressing cells) and the feature plot of RIPOR2 expression confirm that MEK1ca cell-autonomously activates RIPOR2 in transfected cells (Fig. 2 F-G). The expression pattern of RIPOR2 had not been described in the chicken embryo. We have shown by *in situ* hybridization that *RIPOR2* is not expressed in the neural tube or more generally in the trunk of two- and three-day-old chicken embryos (Fig. 2H). Its expression in the head placodes and at later stages is consistent with its described function in controlling otic vesicle and muscle development in vertebrates (Fig. 2H and Supplementary Fig. 3) ^46,47^. We found that it is also expressed in the intermediate zone of the differentiated neural tube from E4, as well as in the dorsal root ganglia (Supplementary Fig. 3). We confirmed by *in situ* hybridization in tissue sections after *in ovo* electroporation that overactivation of ERK1/2 by MEK1ca expression leads to ectopic expression of *RIPOR2* in the trunk neural tube of the chicken embryo (Fig. 2I-J and Supplementary Fig. 4). MEK1ca-induced ectopic expression of *RIPOR2* one day after electroporation was observed in all transfected cells, regardless of the dorsoventral zone of the neural tube (Fig. 2I), including melanocyte precursors located in the most dorsal part of the neural tube.

Altogether, these data show that the *RIPOR2* gene is a general and conserved transcriptional target of the RAS/ERK oncogenic signaling pathway, including in melanocyte progenitors, and is a good candidate for inducing multinucleation when abnormally expressed.

### *In vivo* gain-of-function of RIPOR2 in the chicken embryo trunk neural tube promotes multinucleation

To understand the consequences of ectopic RIPOR2 expression in an *in vivo* context, we utilized trunk neural tube development in the chicken embryo model and performed a RIPOR2 gain-of-function experiment at E2, the stage of MEK1ca transfection ^22^. At this stage, RIPOR2 is not endogenously expressed at the electroporation site (Fig. 2H). RIPOR2 has two major isoforms (iso1 and iso2, with 1068 and 591 amino acids in humans) ^43^. We focused on the shorter iso2, which has a sequence entirely included in the longer isoform and shows stronger affinity to RHOA ^43^. We performed a RIPOR2 gain-of-function experiment on two-day-old chicken embryos by electroporating a vector (pCAGGS-RIPOR2-iresGFP) co-expressing wild-type chicken RIPOR2 iso2 (602 aa) and GFP as a reporter into the right side of the trunk neural tube. This resulted in neuroepithelial disorganization observed as early as one day after electroporation, with cells invading the neural tube lumen (Supplementary Figs. 5 and 6). Morphological changes in the neural tube are visible both in whole-mounted embryos using binocular fluorescence microscopy (Supplementary Fig. 5B, n=22 embryos, phenotype observed in 100% of the embryos one day after electroporation; Supplementary Fig. 5 D, n=19 embryos, in 100% of embryos two days after electroporation) and in tissue sections with F-ACTIN staining (Supplementary Fig. 5C,E). The controlateral (non-electroporated) side of the neural tube, or the neural tube electroporated with a control vector (pCAGGS) expressing only GFP (n>20 embryos), exhibits normal neuroepithelial organization (Supplementary Fig. 6). Immunofluorescence in tissue sections using antibodies against the progenitor marker SOX2 and the pan-neuronal marker TUJ1 reveals neuroepithelial disorganization induced by RIPOR2 electroporation (Supplementary Fig. 7), and TUJ1 staining was observed in cells at the apical face of the neural tube.

Immunofluorescence staining of tissue sections one and two days after electroporation with an anti-cleaved-caspase 3 (CASP3) antibody reveals that RIPOR2 gain-of-function in the trunk neural tube triggers massive apoptosis in electroporated cells (Fig. 3A and Supplementary Fig. 8). Interestingly, two days after electroporation, we observed that some CASP3 positive cells were multinucleated (Fig. 3A) (n=3, observed in 3/3 embryos and not observed in the control condition). To decrease cell death, we co-electroporated a vector expressing RIPOR2 into the trunk neural tube of chicken embryos along with a vector expressing the P35 protein, known to inhibit apoptosis ^48^. In this context, we observed multinucleated electroporated cells in all embryos analyzed three days after electroporation with RIPOR2 and P35 condition (100%, n=4, Fig. 3B-C); this was not observed in control embryos electroporated with P35 alone (0%, n=3, Supplementary Fig. 9). F-ACTIN staining reveals that, similar to RIPOR2 electroporation without P35, co-transfected cells accumulate at the apical side of the neuroepithelium, loss their apico-basal polarity and invade the lumen, and are positive for TUJ1 staining (Fig. 3B-C and Supplementary Fig. 10). We also observed cells with ectopic protrusions, which is consistent with findings from a RIPOR2 gain-of-function study in cell culture (Fig. 3C) ^42^.

**Figure 3:**
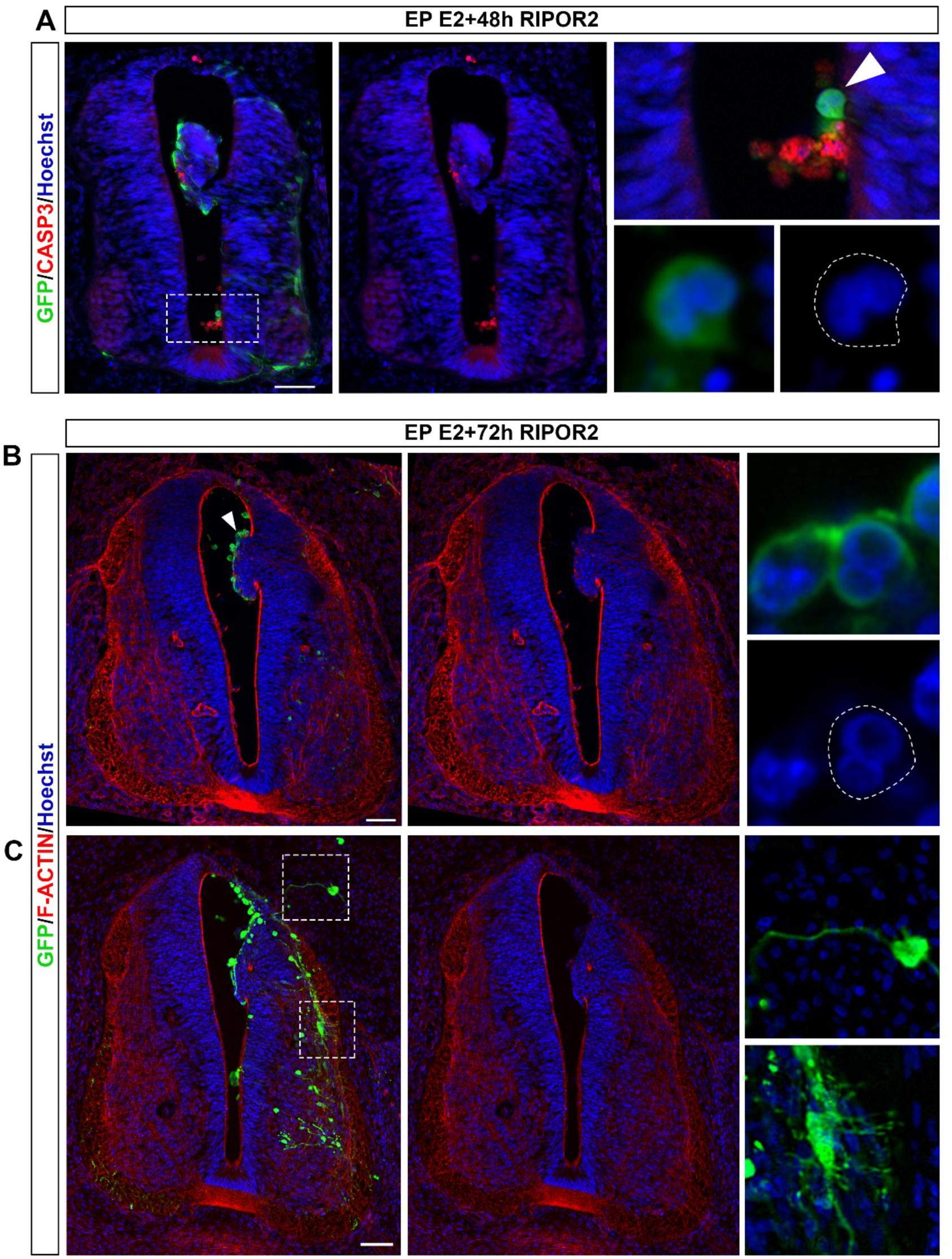
The gain of function of RIPOR2 in the trunk neural tube of chicken embryo promotes cell multinucleation. **A-** Immunofluorescence with anti-GFP and anti-CASP3 antibodies on trunk transverse section of chicken embryo two days after electroporation of RIPOR2. RIPOR2 gain of function induces cells death, but also the apparition of cells with several nuclei (white arrowhead) **B-C-** Immunofluorescences with anti-GFP antibody and F-ACTIN staining on trunk transverse sections of chicken embryo three days after co-electroporation of RIPOR2 + P35 vectors, **B**- which promotes the multinucleation of cells (white arrowhead) and **C**- and ectopic cellular protrusions (dotted boxes). Blue is Hoechst staining in all the panels. Scale bar: 50µm

Overall, these results demonstrate that in an *in vivo* developmental context, RIPOR2 promotes multinucleation when ectopically expressed in a tissue consisting exclusively of mononucleated cells.

### RIPOR2 is expressed in benign human melanocytic nevi and melanomas, but not in healthy skin

We then asked whether the RIPOR2 gene might be relevant in human cutaneous melanoma, specifically, whether its expression was ectopically induced by RAS/ERK oncogenic activity in human melanoma cells, particularly early in disease progression.

To determine whether RIPOR2 is ectopically expressed in melanocytes of human melanoma early in the disease, we performed immunofluorescence with a RIPOR2 antibody on biopsies of healthy human skin tissue (Fig. 4), of benign melanocytic nevi, and of intermediate melanocytic neoplasms (morphologically between dysplastic nevus and melanoma *in situ*) (Fig. 5, Supplementary Figs. 11-13, and Supplementary Table 2). To validate RIPOR2 antibody, we performed immunofluorescence with a RIPOR2 antibody on transverse sections of human placenta at 10 weeks of gestation to validate the specificity of the antibody. Staining with the RIPOR2 antibody is localized to the outer edge of the syncytiotrophoblast ^41^ and to immune cells ^43,44^, as previously described (Supplementary Fig. 11A). Then, we performed hematoxylin and eosin staining and immunostaining on adjacent sections of skin biopsies with anti- BRaf^V600E^ and SOX10 (melanocyte marker) antibodies. We found that although RIPOR2 is not expressed in melanocytes of healthy skin (Fig. 4), it is expressed in the cytoplasm of melanocytes in benign melanocytic nevi (Supplementary Fig. 11B-D and Supplementary Fig. 12) and intermediate melanocytic lesions (Fig. 5 and Supplementary Fig. 13). In intermediate melanocytic lesions, RIPOR2 is expressed in a pattern similar to that of BRaf^V600E^-positive cells (nested pattern) in SOX10-positive areas (Fig. 5). As expected, RIPOR2 is also expressed in immune cells of the skin (Fig. 5 and Supplementary Fig. 13). In nine analyzed skin lesion biopsies, we observed ectopic expression of RIPOR2 in seven, suggesting that this is a common phenomenon in nevi and melanoma (Supplementary Table 2).

**Figure 4:**
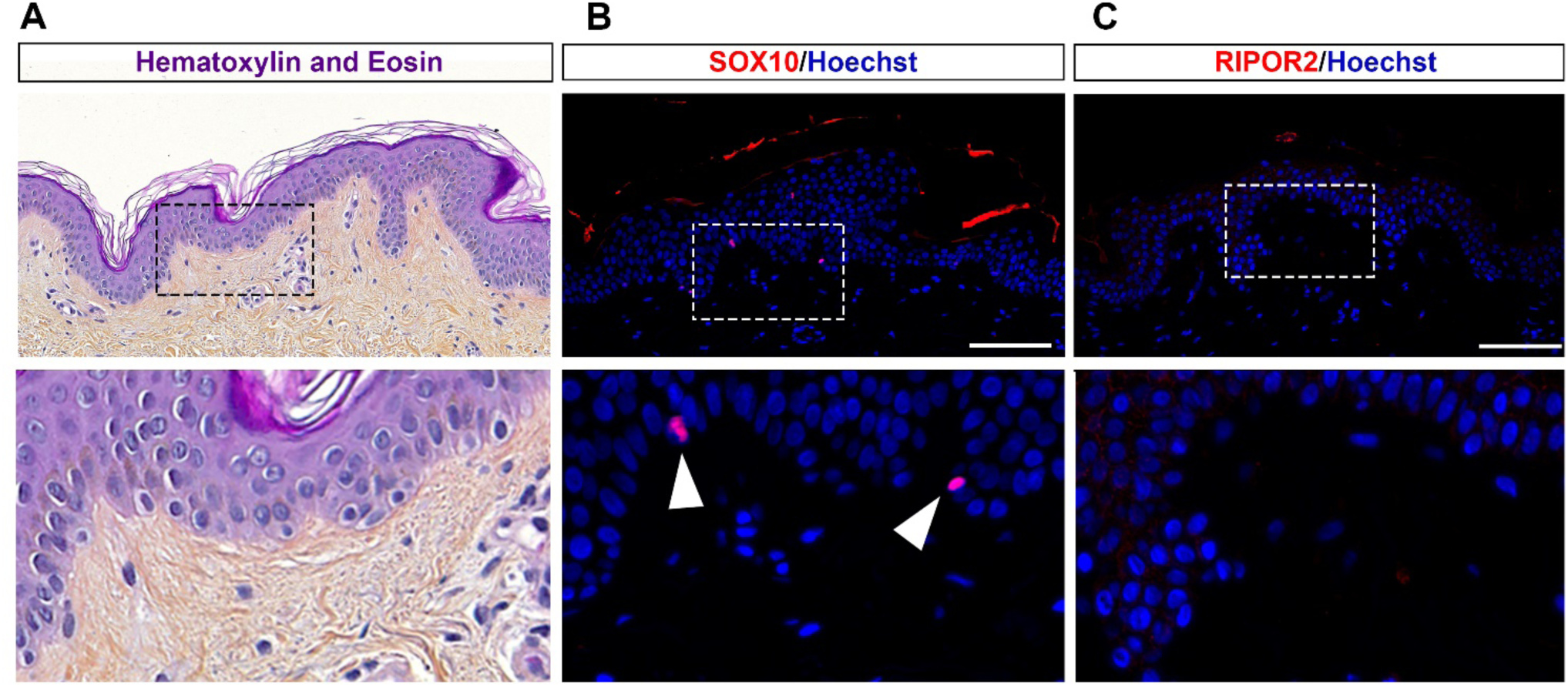
RIPOR2 is not expressed in healthy human epidermal melanocytes. Adjacent sections of healthy skin at the border of an early melanoma biopsy. Dotted boxes are magnified in the bottom panel **A-** Haematoxylin and eosin stain (H&E stain). **B-** Immunofluorescence with an anti-SOX10 antibody shows melanocytes scattered in the epidermis (white arrowhead) **C-** Immunofluorescence with an anti-RIPOR2 antibody shows that it is not expressed in the epidermis including in melanocytes. Blue is Hoechst staining in all the panels. Scale bar: 100µm

**Figure 5:**
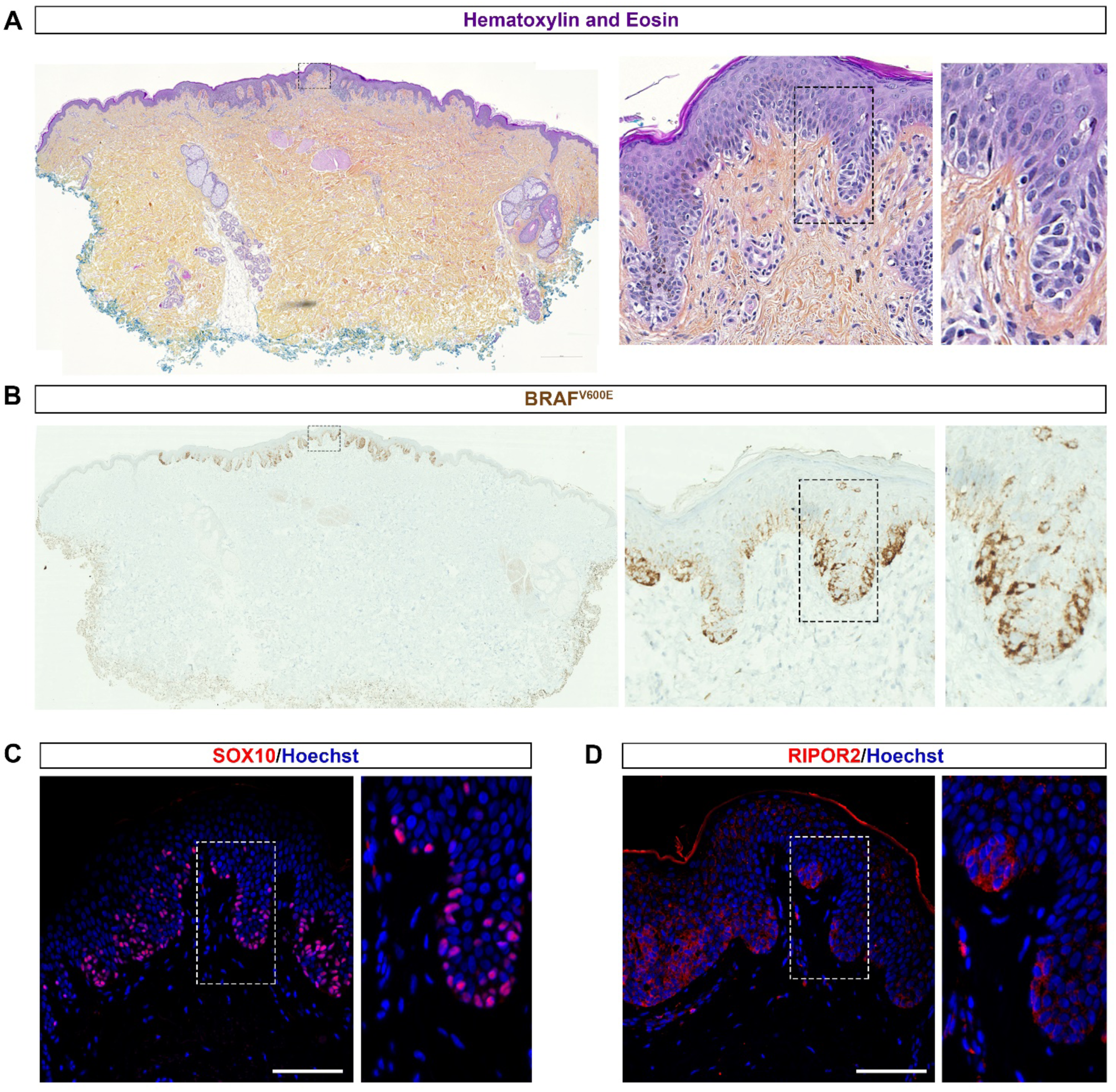
RIPOR2 is expressed in the melanocytes of an intermediate-grade melanocytic lesion. Adjacent sections of an early BRAF^V600E^-positive intraepidermal melanoma. Dotted boxes are magnified in the adjacent panels. **A-** H&E stain shows tissue disorganization in the centre of the section. **B-**Immunohistochemistry with an anti-BRAF^V600E^ antibody stains the mutated melanocytes and highlights the malignant lesion zone. **C-**Immunofluorescence with anti- SOX10 and **D-** anti-RIPOR2 antibodies demonstrated that RIPOR2 is expressed in SOX10+epidermal nests of BRAF^V600E^-positive melanocytes, suggesting that it is expressed in malignant melanocytes. The RIPOR2 staining is cytoplasmic. Blue corresponds to Hoechst nuclear staining in all panels. Scale bar: 100µm.

Bioinformatics data confirm that RIPOR2 is expressed in melanocytes and immune cells of human melanoma. Data mining analysis using BBrowser2 ^49^ with single-cell RNAseq data from Tirosh et al. (2016) ^50^ indicates that RIPOR2 expression in melanocytes of human melanoma is a characteristic feature of most melanomas. Indeed, single-cell RNA analysis of 19 melanoma samples, using violin plots on melanocyte cells, showed that despite some variation among patients, 18 of 19 samples express RIPOR2 in melanocytes (Supplementary Fig. 14 A-E). RIPOR2 is highly expressed in blood/immune cells, compared to melanocyte cells, with little variation among patients (Supplementary Fig. 15 A-C).

Altogether, immunohistochemistry of human skin samples and single-cell RNAseq data of human melanoma show that ectopic expression of RIPOR2 in melanocytes in pre-cancerous lesions, intermediate melanoma, and late-stage melanoma is a common event.

### RIPOR2 is expressed in human melanoma cell lines and its expression is regulated by the RAS/ERK pathway

Since RIPOR2 is expressed in transformed human melanocytes, we next investigated whether its expression was dependent on the RAS/ERK pathway in these cells. By immunofluorescence, we found that RIPOR2 is expressed in the cytoplasm of melanoma cells (SK-MEL-28), exhibiting high expression in approximately 5% of SK-MEL-28 cells. RIPOR2 expression in SK-MEL-28 cells depends on the ERK pathway, as 24-hour treatment with the ERK inhibitor SCH772984 (Fig. 6B) decreased RIPOR2 expression (Fig. 6A, C, and D). By data mining an RNAseq dataset from the human melanoma cell line A375^51^, we found that the transcriptomic expression of RIPOR2 in these cells also depends on the RAS/ERK pathway. Indeed, after three hours of incubation with the ERK inhibitor SCH772984, the MEK inhibitor PD0325901 or the BRAF inhibitor vemurafenib, RIPOR2 expression was significantly reduced (Fig. 6E).

**Figure 6:**
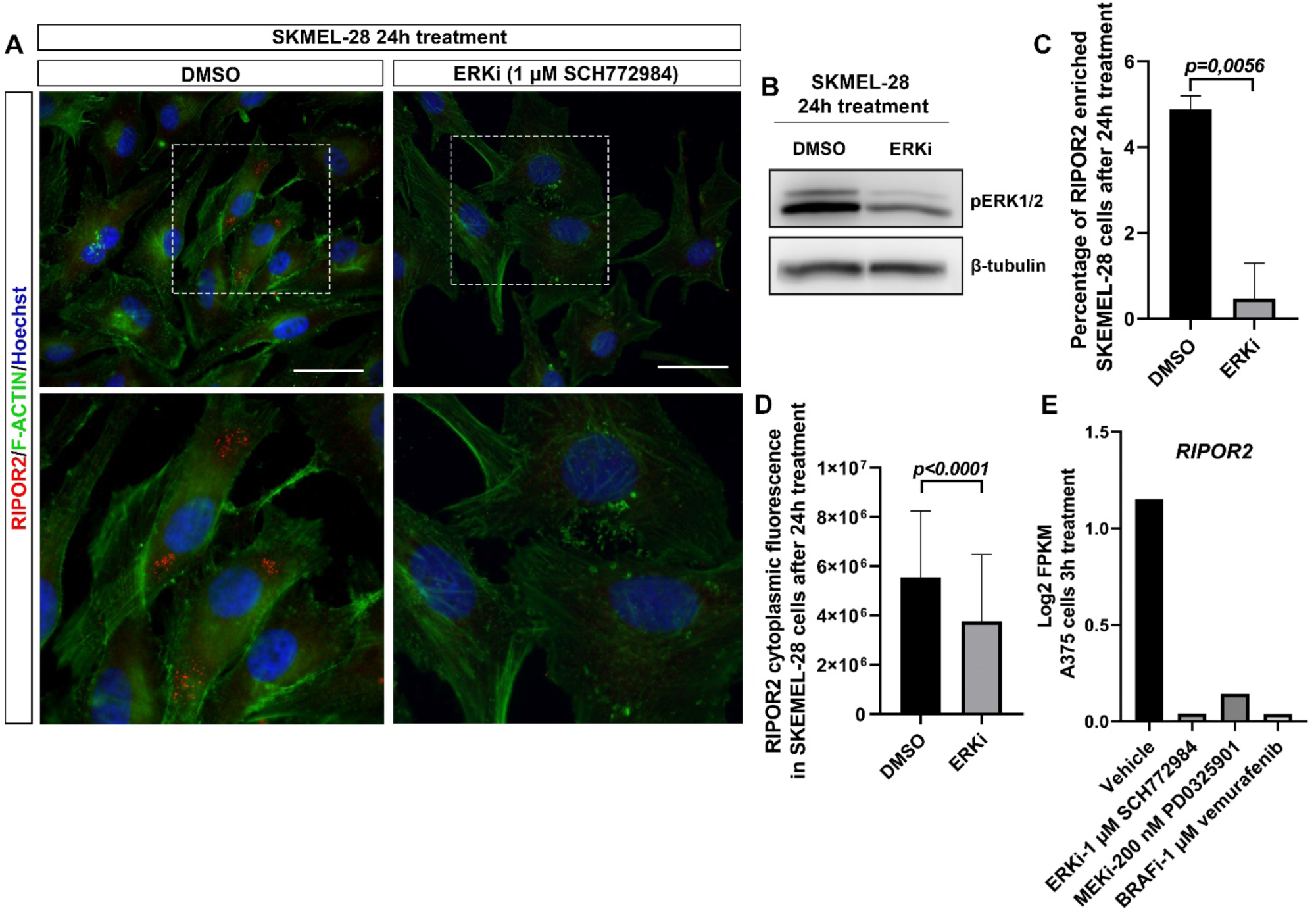
*RIPOR2* expression in human melanoma cell lines is dependent of ERK1/2 activity. **A-** Immunofluorescence with an anti-RIPOR2 antibody and F-ACTIN staining in SKMEL-28 cell line, treated either with DMSO (control) or ERK inhibitor (ERKi - SCH772984) for 24h. In the control condition, RIPOR2 protein is expressed in the cytoplasm and is enriched in a few cells. White dotted boxes are magnified in the bottom panel. ERKi treatment downregulates RIPOR2 expression. Blue is Hoechst staining. Scale bar: 50µm. **B-** Confirmation by western blot that ERKi treated cells display a downregulation of pERK1/2. **C-** ERKi treatment leads to a diminution of the number of SKMEL-28 cells with an enriched expression of RIPOR2 **D-** In these cells, the cytoplasmic corrected cell fluorescence was measured (DMSO, 247 cells; ERKi, 219 cells of 3 independent experiments; two-tailed Mann–Whitney test, error bars represent s.d) which demonstrated global downregulation of RIPOR2 expression. **E**- RNAseq data from Yue et al., 2017 show the FPKM (Fragments per kilo base per million mapped reads) level of *RIPOR2* transcripts in the A375 melanoma cell line, stimulated for 20 minutes with EGF and then incubated 3 hours with either vehicle, ERKi (1µM SCH772984), MEKi (200 nM PD0325901), or BRAFi (1µM vemurafenib). This highlights that RIPOR2 is also expressed in A375 cells and that its transcriptomic expression is also dependent on the RAS-ERK pathway.

Overall, our data demonstrated that the transcriptional expression of RIPOR2 depends on the RAS/ERK pathway in melanoma cells. Ectopic expression of RIPOR2 in human melanocytic lesions could therefore be the consequence of overactivation of RAS/ERK triggered by a driver mutation such as BRaf^V600E^.

### RIPOR2 promotes multinucleation in human melanoma cell line

Since we have shown that RIPOR2 is ectopically expressed in human melanocytic lesions and promotes ectopic multinucleation *in vivo* in the chicken embryo neural tube, we next sought to test whether RIPOR2 also promotes multinucleation in human melanoma cells. We therefore tested whether RIPOR2 gain-of-function increases the number of multinucleated cells in human melanoma cells. Transfection of SK-MEL-28 melanoma cells with a vector expressing human RIPOR2 (iso2) fused with GFP at the C-terminus ^43^ induces an increase in the number of multinucleated cells 48 hours after transfection (8.4% in cells transfected with a vector expressing only GFP versus 26.2% in cells transfected with the vector expressing hRIPOR2- GFP, Fig. 7). This phenotype is not limited to melanoma cells, as transfection of hRIPOR2- GFP into HeLa cells in both transiently transfected (Supplementary Fig. 16) and stable inducible cell lines (Supplementary Fig. 17) also induced an increase in the number of multinucleated cells. This increase was observed as early as one day after transfection or following doxycycline induction (3.6% in cells transfected with a vector expressing only GFP versus 12.3% in cells transfected with the vector expressing hRIPOR2-GFP, Supplementary Fig. 16; 3.3% in cells expressing only GFP versus 6.8% in cells expressing hRIPOR2-GFP, Supplementary Fig. 17). We conclude that RIPOR2 promotes multinucleation in human cancer cell lines, including melanoma.

**Figure 7:**
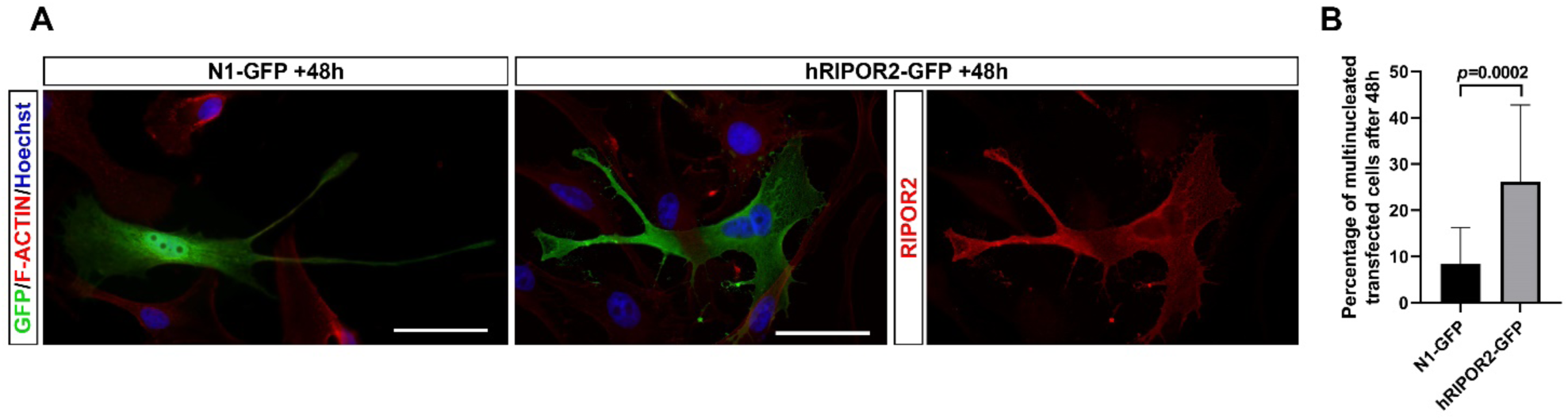
*RIPOR2* overexpression in human melanoma cell line SKMEL-28 promotes multinucleation. **A-** Immunofluorescence with anti GFP, anti- RIPOR2 antibodies and F-ACTIN staining in SKMEL-28 cell line, transfected either with a control plasmid expressing only GFP (N1-GFP) or human RIPOR2-GFP (hRIPOR2-GFP) for 48h. The transitory expression of h RIPOR2 increases the number of transfected (GFP+) multinucleated cells, quantify in **B-** represented as the percentage of transfected multinucleated cells (N1-GFP, 127 cells, h RIPOR2,165 cells, 4 independent experiments, Fisher’s exact test). Blue is Hoechst staining. Scale bar: 50µm

## DISCUSSION

In this study, we identified a mechanism downstream of RAS/ERK oncogenic activation that may contribute to the transformation of benign nevi into melanoma. Although the risk may be very low ^7^, this implies that overactivation of RAS/ERK signaling is not sufficient to cause tumor transformation. In this study, we combined *in vivo* experiments using the chicken embryo model, bioinformatic analysis, human cell culture experiments, and human skin biopsies. We showed that the RAS/ERK oncogenic pathway triggers ectopic expression of RIPOR2, and that the RIPOR2 protein promotes the emergence of multinucleated cells. There is some evidence that multinucleated melanocytes are a source of tumor-initiating cells^16^. One reason why the transition from nevi to melanoma is rare may be that the appropriate response to multinucleated melanocytes is cell death or senescence ^16^. However, multinucleation-induced aneuploidy may fuel cellular heterogeneity. Inactivation of RIPOR2 may therefore limit cancer aggressiveness by limiting multinucleation. Some individuals develop only a few nevi, while others develop hundreds. The number of nevi is a combination of environmental mutagens, including ultraviolet radiation, and hereditary factors ^52^. Although resection of suspicious nevi is a common and effective strategy to limit the development of melanoma^53^, it is not realistic in patients with hundreds of nevi or with giant congenital melanocytic nevi, and has never been directly tested for prophylaxis. It is therefore important to have alternative prevention solutions for these patients, because once initiated, melanoma lacks satisfactory therapeutic solutions. Since the RAS/ERK pathway is overactivated in almost all types of cancer^54^, RIPOR2 may also have a similar function in other solid tumors where the RAS/ERK pathway is deregulated. Thus, it constitutes a therapeutic target for most cancers.

One mechanism by which RIPOR2 may promote tumor cell multinucleation is by disrupting mitotic spindle formation and causing cytokinesis failure. Indeed, RIPOR2 has been described to control proliferation via its interaction with two proteins involved in cell proliferation, namely the HDAC6 deacetylase and the 14-3-3 scaffolding protein. RIPOR2 expression disrupts mitotic spindle formation, leading to mitotic failure in T-cells and HeLa cells ^55^. Mitotic failure is known to lead primarily to cell death ^56^, as observed after RIPOR2 gain-of-function in T-cells and HeLa cells ^55^. The increased cell death phenotype we observed after RIPOR2 gain-of-function in the chicken embryo trunk neural tube could therefore be a consequence of mitotic failure that triggers apoptosis. In the context of RIPOR2 ectopic expression induced by oncogenic RAS/ERK activation leading to inhibition of cell death ^57^, mitotic failure might escape cell death more frequently than in the gain-of-function of the protein alone.

A second mechanism by which RIPOR2 could promote multinucleation in tumor cells is by inducing cell-cell fusion. Indeed, RIPOR2 is expressed in two developing tissues at the beginning of their multinucleation, the placenta ^58^ and skeletal muscle ^42^, and has been suggested to control myocyte cell fusion ^42^. Its ectopic expression triggered by RAS/ERK overactivation could hijack its developmental function and promote ectopic cell-cell fusion. The fusion of one cell with another occurs in development, upon injury, and cell fusion is suggested to be a possible cause of some cancers, as it could explain the occurrence of multiple genetic changes considered to underlie cancer. Indeed, cell fusion likely stimulates tumor evolution by compromising chromosome and DNA stability and/or by generating phenotypic diversity; however, it remains unproven *in vivo* whether a cell fusion event can initiate malignancy and direct tumor evolution. Some have reported fusion events giving rise to tumor cells with CSC characteristics, e.g., in the spontaneous fusion of two human breast cancer cell lines ^59^, two transformed human fibroblast cell lines ^60^, or mesenchymal adipose-derived stem cells with breast cancer cells ^61^. In all these cases, the hybrid cells expressed more stem cell markers associated with higher tumorigenic potential. In addition, artificial cell fusion events triggered by polyethylene glycol in normal, non-transformed, cytogenetically stable epithelial cells can initiate chromosomal instability, DNA damage, cellular transformation, and malignancy ^62^. Clonal analysis of fused cells reveals that the karyotypic and phenotypic potential of tumors formed by cell fusion is established immediately or within a few cell divisions after the fusion event, without further ongoing genetic and phenotypic plasticity. The subsequent evolution of these tumors reflects selection from the initial diverse population rather than ongoing plasticity of the progeny ^62^. This suggests that a cell fusion event could both initiate malignancy and fuel subsequent tumor evolution.

In conclusion, it is likely that RIPOR2-induced multinucleation impacts the aggressiveness of melanocytic lesions and other tumor cells in which it is ectopically expressed downstream of RAS/ERK overactivation. It would therefore be relevant to evaluate its potential as a therapeutic target in early melanoma and other cancers. Further studies could also identify the cellular and molecular mechanisms by which RIPOR2 promotes multinucleation in tumor cells downstream of the RAS/ERK pathway.

## MATERIALS AND METHODS

### Ethics statement

The chicken embryos analyzed were in early stages of embryonic development (between E2 and E5). Therefore, no specific approval from the Institutional Animal Care and Use Committee was sought (French decree 2013-118 from 1^st^ February 2013 and Directive 2010/63/EU (http://data.europa.eu/eli/dir/2010/63/2019-06-26) of the European Parliament and of the Council of 22 September 2010 on the protection of animals used for scientific purposes).

Unstained slides, cut from archival FFPE material of patients with melanoma, melanocytic nevi, and dysplastic nevi were collected from a previous study, and for which all patients gave informed consent to include their anonymized samples in the APHM Biobank ^63^. The slides were provided by the Assistance Publique Hôpitaux de Marseille, Biological Resources Center (BRC AP-HM Biobank), CRB-TBM component (NF S96-900 & ISO 9001 v2015 Certification), from the dermatopathology collection of Pr. Gaudy-Marqueste and Dr. Nicolas Macagno. These bioresources belong to a biological sample collection declared to the French Ministry of Health (Declaration: DC-2013-1781) whose use for research purposes was authorized by the French Ministry of Higher Education, Research and Innovation (Authorization: AC-2011-2018-3105). Adjacent unaffected skin from the same patients was also used as controls.

Human embryonic material was obtained after informed consent through the HuDeCA INSERM Transverse Program under Biomedicine Agency protocol PFS14-011 and ministerial authorization DC-2019-3716.

### Chicken embryos

Fertilized chicken eggs were obtained from EARL les Bruyeres (Dangers, France) and incubated horizontally at 38°C in a humidified incubator. Embryos were staged according to the developmental table of Hamburger and Hamilton (HH) ^64^ or by days of incubation (E).

### *In ovo* electroporation and plasmids

Uni- or bilateral neural tube *in ovo* electroporation was performed around HH12, as already described ^65^. The plasmids used co-express cytoplasmic or nuclear GFP (pCAGGS or pCIG, respectively) and the coding sequence (CDS) of the gene of interest. The pCAGGS and pCIG plasmids were used alone as controls. Vectors used were: pCIG, pCIG-MEK1ca ^66^, pCAGGS, pCAGGS-RIPOR2 (co-expressing GFP and the coding sequence (CDS) of the gene of isoform 2 of chicken RIPOR2) and pCAGGS-P35 (co-expressing GFP and the CDS of the P35 protein^48^). The plasmids used for electroporation were purified using the Nucleobond Xtra Midi kit (Macherey-Nagel). The final concentration of DNA delivered to each embryo for electroporation was up to 2 µg/µl.

### Single-nuclei preparation and 5′ Gene expression single nuclei library preparation

For each of the two conditions (pCIG or pCIG-MEK1ca), 24 trunks of embryos one day after electroporation were pooled, frozen in liquid nitrogen, and stored at – 80°C in subgroups of six samples each. Single nuclei were isolated using an in-house protocol developed by the CYBIO core facility at Institut Cochin, Paris (previously described ^67^). Samples were then suspended in 2% BSA in PBS, filtered through 100 µm cell strainers (VWR), centrifuged twice for 10 min at 500g, and resuspended in 2% BSA in PBS. Nuclei were incubated with an Alexa Fluor® 647 anti-Nuclear Pore Complex Proteins Antibody Mab414 (BioLegend), then sorted using a FACSAria III (BD Biosciences) with the 85 μm nozzle and the BD FACSDIVA software. Sorted nuclei were immediately processed on a Chromium Controller (10x Genomics). The mRNA was reverse-transcribed, converted to barcoded cDNA with sample indexing, purified using DynaBeads, and amplified by PCR. To construct the 5′ gene expression library, the amplified barcoded cDNA was fragmented, end-repaired, poly-A-tailed, sample-indexed, and double-size selected with SPRI beads (average size 450 bp). The DNA was quantified, and fragment size distribution of the libraries was determined using the Qubit dsDNA HS assay kit (ThermoFisher, Q32851) and Agilent 2100 BioAnalyzer High Sensitivity DNA kit (Agilent Technologies, 5067-4626). Pooled libraries were then sequenced on an Illumina Nextseq 500 sequencing platform to a minimum sequencing depth of 20,000 reads per nucleus using read lengths of 26 bp read 1, 10 bp dual indexes, and 90 bp read 2.

### Bioinformatics

Fastq files were then aligned, counted, and assigned to nuclei using the ‘CellRanger’ algorithm (version 6.0.1, with STAR v2.7.2a), based on the Ensembl GRCg6a_v6 reference. The quality controls were performed with R (version 3.6.3) and the Seurat package (seurat_4.0) on raw unfiltered expression matrices. We kept cells that detect more than 200 genes and less than 2500 genes and that detect less than 5% of mitochondrial genes.

After filtering, we took both expression matrices (PCIG-Bis and MEK1ca samples) and checked if we have a batch effect (ie, technical variations that generate a bias, preventing direct comparison between several samples). To do this, we normalized data together with the function “NormalizeData” from seurat (v5.1.0, R v4.3.1), with the method “LogNormalize”. All parameters were set to default except for the “scale.factor” where we took the median of all total counts. Then we performed a Principal Component Analysis (PCA) with the RunPCA function (default parameters). When inspecting the PCA, we see a good overlap between the two samples confirming the fact that there were no technical variations and no need to correct a potential batch effect. We cluster nuclei thanks to FindNeighbors and FindClusters functions. We took the first 50 first Principal Components (PCs) based on the results of the JackStraw methods implemented in Seurat where we saw that all PCs got a p-value inferior to 1e-30, and we looked at partitions with different resolutions (0.2 to 1.2 with a step of 0.2). We inspect clustering results with the R package clustree (v0.5.1). It represents the relationships and the distribution of the cells within the clusters at different resolutions, when a cluster has several clusters of origin at a lower resolution it probably means that we took too high a resolution. Since the results were quite clean, we took a resolution of 0.6 because it seemed to be a good compromise between a fine clustering without over-clustering. After, we export the results to a cloupe file for further analysis in Loupe Browser thanks to the R package loupeR (v1.1.1). Feature plots, violin plots, and Differential Expression were done using Loupe Browser (8.0.0). “Advanced Selection” of Loupe Browser was used to make the custom group expressing GFP>1 for MEK1ca and Control conditions. Single-cell RNAseq data mining on human melanoma ^50^ (t-SNE (visualization of cell types among patients), feature plots and violin plots), was done using BBrowser2 (https://bioturing.com/bbrowser/download)^49^.

### Immunofluorescence on tissue section

Tissue preparation, sections and immunofluorescence were performed as already described ^22^. The following primary antibodies were used in this study: chicken anti-GFP 1:1000 (1020 AVES), rabbit anti-SOX2 1:500 (AB5603 Merck Millipore), mouse anti-Tuj1 1:500 (801202 Biolegend), rabbit anti-Caspase 3 1:500 (Asp175, CST 9661). The secondary antibodies used were: anti-chicken, anti-rabbit, anti-mouse, or anti-rat with fluorochromes (488, 568, or 647) at 1:500 (Alexa Fluor, abcam). They were incubated for one hour in the blocking solution containing Hoechst (1:1000). F-ACTIN staining was performed using Phalloidin- AlexaFluor568 (1:70) (ThermoFisher). Slides were mounted in ThermoFisher Shandon Immu- Mount and imaged with a Zeiss Z1 Apotome or a Zeiss LSM 780 confocal microscope.

### Immunofluorescence on paraffine sections

The sections were deparaffinized in xylene twice for 5 min, rehydrate with sequential washes of 100%, 96%, 70%, 50% EtOH, and then rinsed with distilled water. Antigens were retrieved in a boiling solution of Antigen Unmasking Solution (pH 6, Vector H-3300) in distilled water. To reduce nonspecific staining, the slides were incubated 20 min in a solution of 50 mM glycine, 0.1 M NH_4_Cl in H_2_O. After blocking for one hour in 2% FBS in PBS with 0.1% Tween- 20, slides were incubated overnight with the primary antibodies in the blocking solution. The primary antibodies used were: rabbit anti-RIPOR2 (1:100, 17015-1-AP, Proteintech) and rabbit anti-SOX10 (1:250, abcam). The next day, slides were washed 5 times in PBS with 0.1% Tween-20, and incubated 2 hours with the secondary antibodies (rabbit anti- fluorochromes 568 or 647 at 1:500, Alexa Fluor, abcam) in the blocking solution, containing Hoechst (1:1000).

### In situ hybridization

The *IL1R1* and *RIPOR2* probes were produced from PCR products amplified from cDNA from the neural tube of an E3 chicken embryo transfected with MEK1ca ^22^ (IL1R1 primers: fw tgccgataaccacagagaga, rev: TAATACGACTCACTATAGGGCccggtctcatcttcagtgga; RIPOR2 primers: fw CGACCTGCCTTATGAAGACC, rev: TAATACGACTCACTATAGGGTCCAGATGCATCACTTCCTG, containing the T7 RNA polymerase promoter sequence). Fluorescent *in situ* hybridization on tissue sections and in whole mounts was performed as described before ^22^.

### Cell culture

HeLa and SK-MEL-28 cell lines were cultured at 37 °C with 5% CO2 in Dulbecco’s Modified Eagle’s Medium (DMEM) GlutaMAX. For the SKMEL-28 cell line, the medium was supplemented with 1% FBS (ATCC) and 1% Gibco Sodium Pyruvate (100 mM).

#### SKMEL-28 ERK inhibitor treatment

The day prior, 4×10^5^ cells/ml were seeded on coverslips in 24-well plates for immunofluorescence and in 100 mm cell culture dishes for Western blotting. The cell medium was supplemented for 24 hours with either SCH772984 (HY-50846) at 1 µM or DMSO (Sigma) at 1:10,000, replaced 3 times within 24 hours.

#### Western blotting

On ice, cells were washed twice with PBS and incubated with RIPA buffer (0.15 M NaCl, 0.01 M Na3PO4 pH 7.2, 2 mM EDTA, 50 mM NaF, supplemented with 0.2 mM Na3VO4 and protease inhibitor (Roche)) for 10 min. Cells were scraped and centrifuged at 15,000 rpm for 10 min. The eluted proteins were heated for 5 min at 95°C and mixed with 6x reducing Laemmli buffer. They were resolved on a 12% SDS-PAGE acrylamide gel and subjected to immunoblotting. After one hour of blocking in 5% BSA in PBS with 0.1% Tween-20, membranes were incubated overnight at 4°C in the blocking buffer containing the following primary antibodies: rabbit anti-Phospho-p44/42 MAPK (ERK1/2) (Thr202/Tyr204) Antibody #9101 (Cell Signaling Technology, 1:1000) and rat anti-β-tubulin (ab15568, Abcam, 1:2500). After 3 washes in PBS with 0.1% Tween-20, the following secondary antibodies were used for 1 hour at room temperature: rabbit anti-HRP and rat anti-HRP (Jackson Immuno, 1:20,000). The kit ECL Western Blot Substrates (ThermoFisher) was used for detection and we used a ChemiDoc (Bio-Rad) for imaging.

#### Transient cell transfection and immunofluorescence

The day prior, 8×10^5^ SK-MEL-28 cells/ml and 6×10^5^ HeLa cells/ml were seeded on coverslips in 24-well plates in complete medium without antibiotics. Transient transfection with the plasmid N1-GFP or N1-hRIPOR2-GFP was performed with X-tremeGENE™ HP DNA Transfection Reagent (Sigma-Aldrich) with 0.5 µg of DNA per well, following the manufacturer’s instructions. Cells were fixed in 4% formaldehyde 4% sucrose in 0.1 M phosphate buffer (0.2 M Na2HPO4, 0.2 M NaH2PO4, 1 M CaCl2) overnight at 4°C, rinsed in 4% sucrose in 0.1 M phosphate buffer, and then rinsed in 15% sucrose in 0.1 M phosphate buffer overnight. Coverslips were then frozen and stored at -20°C until further processing. Coverslips were washed three times in PBS, permeabilized with PBS containing 0.3% Triton, and blocked for 1 hour in 4% BSA and 0.3% Triton in PBS. Primary antibodies used were chicken anti-GFP (1:1000, 1020, AVES) and rabbit anti-RIPOR2 (1:500, 17015-1-AP, Proteintech), incubated overnight in the blocking solution. After three PBS washes, the coverslips were incubated with secondary antibodies (1:500, Alexa Fluor, Abcam), Phalloidin AlexaFluor568 (1:70, ThermoFisher), and Hoechst (1:1000). Coverslips were mounted in ProLong™ Gold Antifade Mountant (ThermoFisher) and imaged with a Zeiss M2 microscope equipped with Apotome.

#### Tetracycline-inducible HeLa cell lines and immunofluorescence

The cDNA encoding hRIPOR2 isoform 2 was subcloned into the pLVX-TetOne-Puro lentivector. HeLa inducible cell lines were established using lentiviral transduction strategy with empty pLVX-TetOne-Puro or hRIPOR2-containing pLVX-TetOne-Puro vectors. Transduced cells were then selected with 1 µg/ml puromycin for 3 days. Cells were fixed with 3.7% formaldehyde for 15 min at room temperature. Cells were washed twice in PBS, permeabilized with PBS containing 0.5% Triton X-100 for 5 min, and then blocked for 30 min with PBS containing 0.2% Tween, 1% BSA, and 1% SVF to prevent non-specific staining. Cells were incubated with primary antibodies diluted in PBS with 0.2% Tween-20 at room temperature for 40 min to 1 hour. After washing in PBS with 0.2% Tween-20, cells were incubated for 30 min with secondary antibodies. Following washing in PBS with 0.2% Tween- 20, coverslips were directly mounted using VECTASHIELD® Antifade Mounting Medium with DAPI. Images of immunofluorescence staining were captured with a fluorescent microscope Leica Inverted 6000 using MetaMorph software.

#### Quantifications of the multinucleated cells

The number of nuclei per transfected cell for transient transfection or in all the cells in an area for stable transfection was counted after acquiring random images on the coverslips. The number of cells counted per condition is indicated in the figure legends and repeated independently at least three times. Fisher’s exact test was performed to analyze the proportion of single-nucleated cells vs multinucleated cells, and the percentage of multinucleated cells for each condition was plotted. The *P*-value was considered significant when P<0.05. All P-values are indicated on the graphs. The error bars represent the standard deviation (s.d.).

## Acknowledgements

We thank Muriel Andrieu at the CYBIO platform (Cochin), and in particular Céline Bertholle and Vaarany Karunanithy who performed the single-nucleus preparation and 5′ gene expression single-nucleus library preparation. We thank Franck Letourneur at the GENOM’IC plarform (Cochin), and in particular Benjamin Saintpierre who performed part of the snRNAseq statistical analyses. We thank the Optical Imaging Platform of the IBDM. We thank Thomas Vannier from the CENTURI Multi-Engineering Platform for his advice on the analysis of single-cell RNAseq data using BBrowser2. We thank Héloïse Toraille and Tamaki Kurosawa for critically reading the manuscript. ARTbio was supported by the CNRS, SU, the Institut Français de Bioinformatique (IFB), and a grant from SIRIC CURAMUS. A.W. was awarded Ph.D. fellowships from La Ligue contre le Cancer and the IBDM. We warmly thank La Ligue contre le cancer, Cancéropôle PACA, AMIDEX, ANR, IBDM, LBD, Dev2A, and IBPS for their founding support.

## AUTHOR CONTRIBUTIONS

Conceptualization: M.C.D., A.W. and E.H.; Methodology: M.C.D., A.W. , A.R., H.E., E.H., T.G.; L.Be. ; N.N. and S.M. ; Software: E.H., L.Be., N.N. and T.G.; Investigation: M.C.D., A.W., A.R., N.M., E.H., T.G., L.Be, N.N., C.G., S.M., N.D., L.Bo., N.C. ; Resources : M.C.D.; A.W. E.H., N.M.; C.G.; S.M.; N. D.; H.E.; Validation: M.C.D., A.W., S.M. and E.H.; Visualization: M.C.D., A.W and H.E.; Writing – original draft: M.C.D., A.W. and H.E.; Writing – review and editing: M.C.D., A.W and H.E.; Supervision: M.C.D ; Project administration: M.C.D..; Funding acquisition: M.C.D. ,Y. G. ,D.D.

