## Supplementary Figures for "RIPOR2 promotes multinucleation of melanoma cells downstream of the RAS/ERK oncogenic pathway"

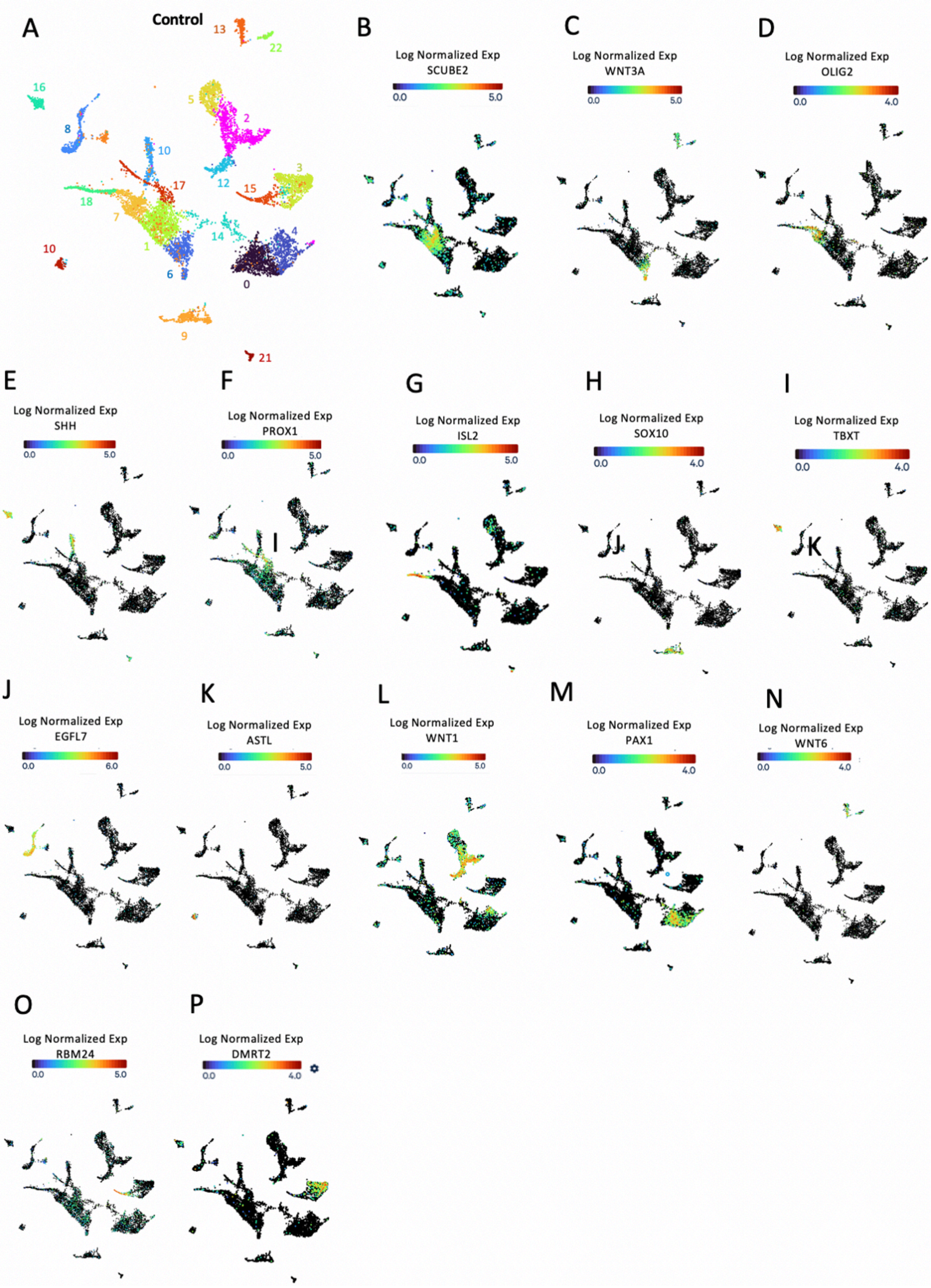

A
Control
13
22
16
5
2
8
10
3
17
12
15
18
7
14
4
1
10
6
0
9
21
B
Log Normalized Exp
SCUBE2
0.0
5.0
C
Log Normalized Exp
WNT3A
0.0
5.0
D
Log Normalized Exp
OLIG2
0.0
4.0
E
Log Normalized Exp
SHH
0.0
5.0
F
Log Normalized Exp
PROX1
0.0
5.0
G
Log Normalized Exp
ISL2
0.0
5.0
H
Log Normalized Exp
SOX10
0.0
4.0
I
Log Normalized Exp
TBXT
0.0
4.0
J
Log Normalized Exp
EGFL7
0.0
6.0
K
Log Normalized Exp
ASTL
0.0
5.0
L
Log Normalized Exp
WNT1
0.0
5.0
M
Log Normalized Exp
PAX1
0.0
4.0
N
Log Normalized Exp
WNT6
0.0
4.0
O
Log Normalized Exp
RBM24
0.0
5.0
P
Log Normalized Exp
DMRT2
0.0
4.0

**Supplementary Figure 1:**

**A -** UMAP plot at snn_res.0.6 for the control condition (pCIG nuclei only). Feature plot of the control condition nuclei only for the SCUBE2 gene (B), the WNT3A gene (C), the OLIG2 gene (D), the SHH gene (E), the PROX1 gene (F), the ISL2 gene (G), the SOX10 gene (H), the TBXT gene (I), the GHFL7 gene (J), the ASTL gene (K), the WNT1 gene (L), the PAX1 gene (M), the WNT6 gene (N), the RBM24 gene (O), and the DMRT2 gene (P).

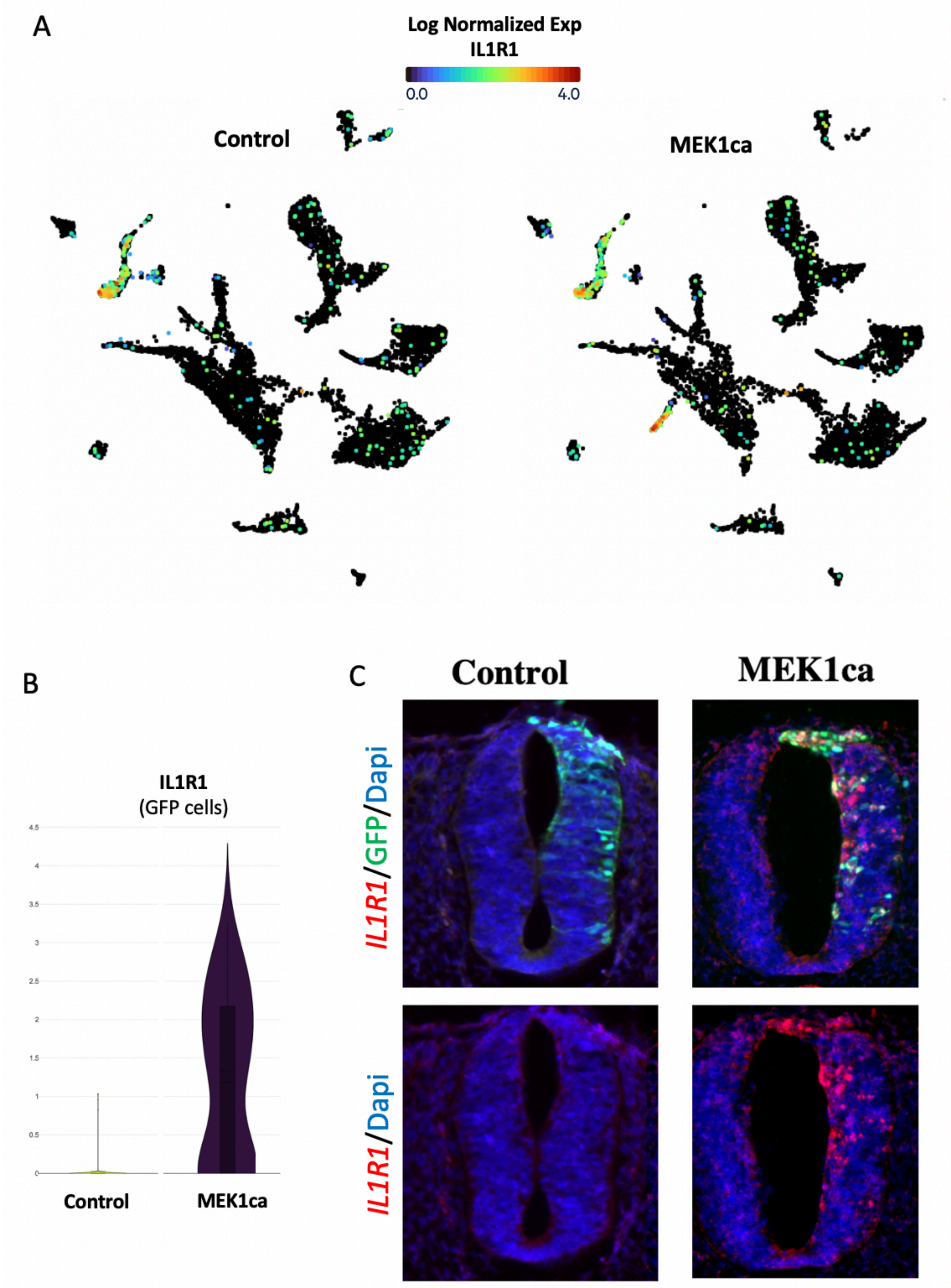
A
Log Normalized Exp
IL1R1
0.0
4.0
Control
MEK1ca
B
IL1R1
(GFP cells)
4.5
4
3.5
3
2.5
2
1.5
1
0.5
0
Control
MEK1ca
C
Control
MEK1ca
IL1R1/GFP/Dapi
IL1R1/Dapi

FIG SUP 2

**Supplementary Figure 2:**

**A -** Feature plot of the two conditions (control and MEK1ca) for the IL1R1 gene. **B -** Violin plot of the IL1R1 in transfected nuclei (GFP > 1). **C -** Fluorescence in situ hybridization with the *IL1R1* probe and immunofluorescence staining with the anti-GFP antibody on transverse trunk sections of chicken embryos one day after electroporation in the control (pCIG) or MEK1ca conditions.

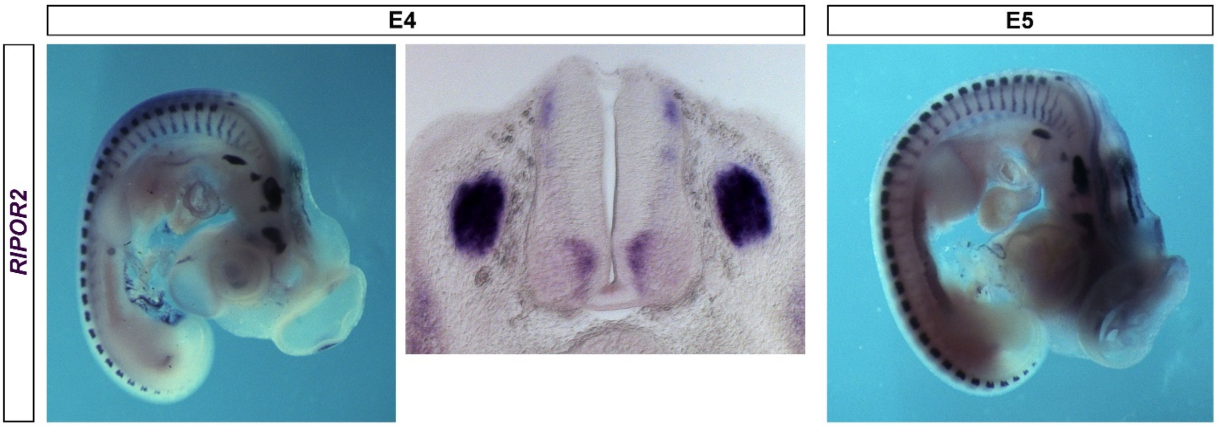


**Supplementary Figure 3:**

Lateral view and transverse section of E4 chicken embryo and lateral view of E5 chicken embryo after whole-mount in situ hybridization with the chicken RIPOR2 probe.

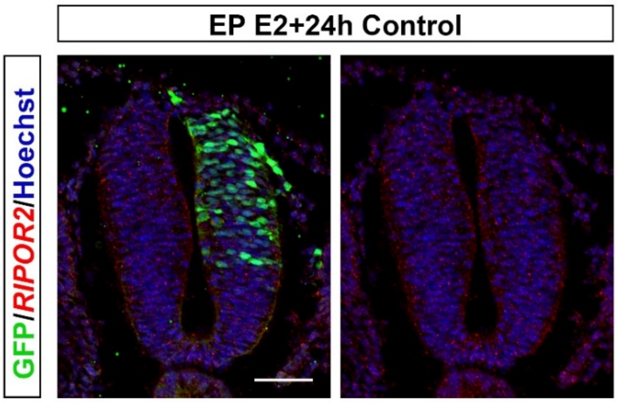


**Supplementary Figure 4:**

Fluorescent *in situ* hybridization with a chicken *RIPOR2* probe and immunofluorescence staining with the anti-GFP antibody on trunk transverse section of chicken embryo one day after electroporation of the control plasmid (pCIG). Blue represents Hoechst nuclear staining. Scale bar: 50 µm

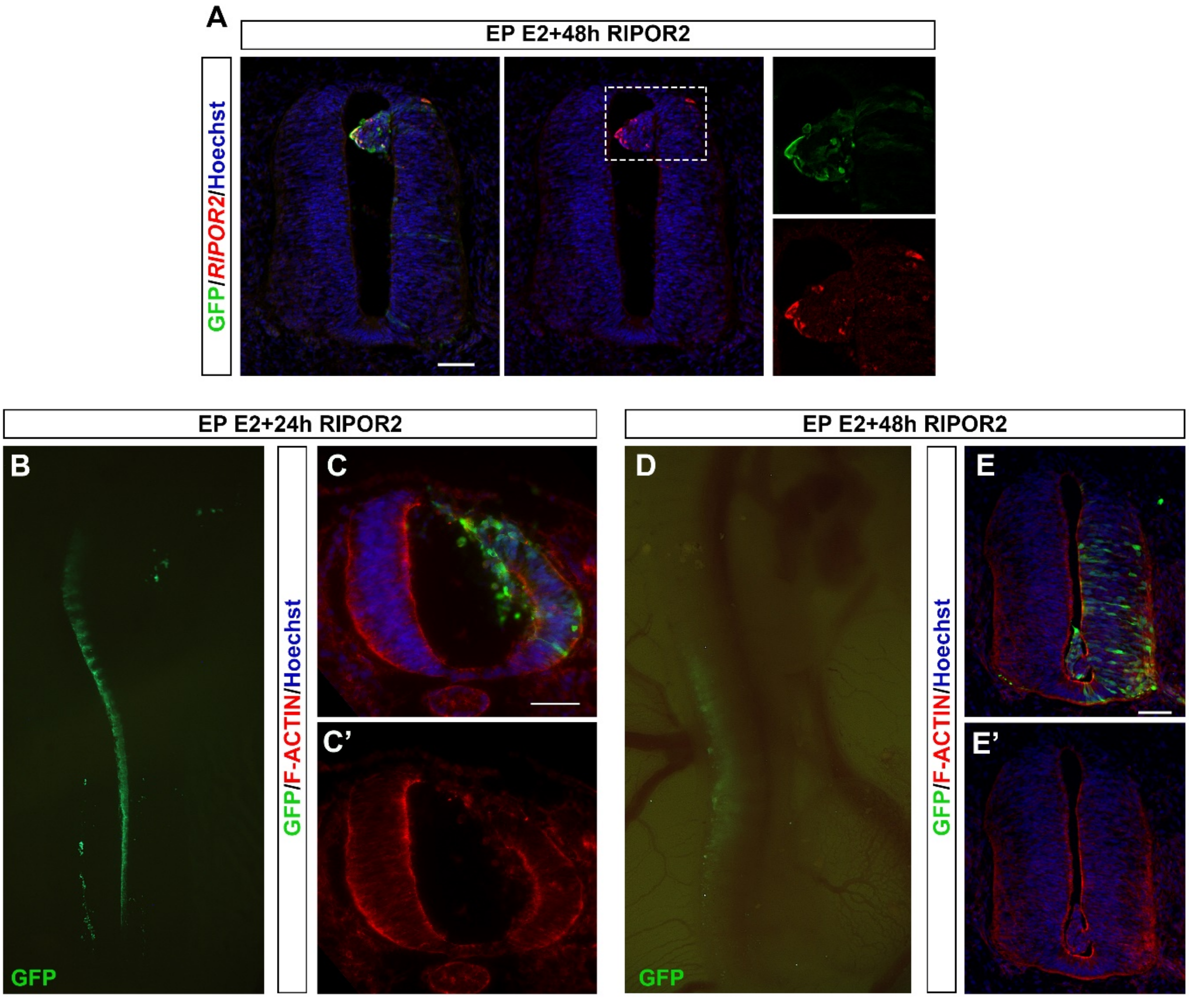


**Supplementary Figure 5: The gain-of-function of RIPOR2 in the trunk neural tube in chicken embryos disrupts the neuroepithelium**

**A -** Fluorescent *in situ* hybridization with a chicken RIPOR2 probe and immunofluorescence with the anti-GFP antibody on trunk transverse section of chicken embryo two days after electroporation of RIPOR2. **B -** Dorsal view of a whole-mount embryo one day after the electroporation of the RIPOR2-expressing vector under fluorescent binocular highlighting GFP+ transfected cells (left side). **C-C' -** Immunofluorescence staining with the anti-GFP antibody and F-ACTIN staining on trunk transverse section of chicken embryo one day after electroporation of RIPOR2, highlighting the disorganization of the neuroepithelium with cells invading the lumen of the neural tube. **D -** Dorsal view of a whole-mount embryo two days after the electroporation of the RIPOR2-expressing vector using a fluorescent binocular microscope. **D-D' -** Immunofluorescence with the anti-GFP antibody and F-ACTIN staining on trunk transverse section of chicken embryo two days after electroporation of RIPOR2. Blue represents Hoechst nuclear staining. Scale bar: 50 µm.

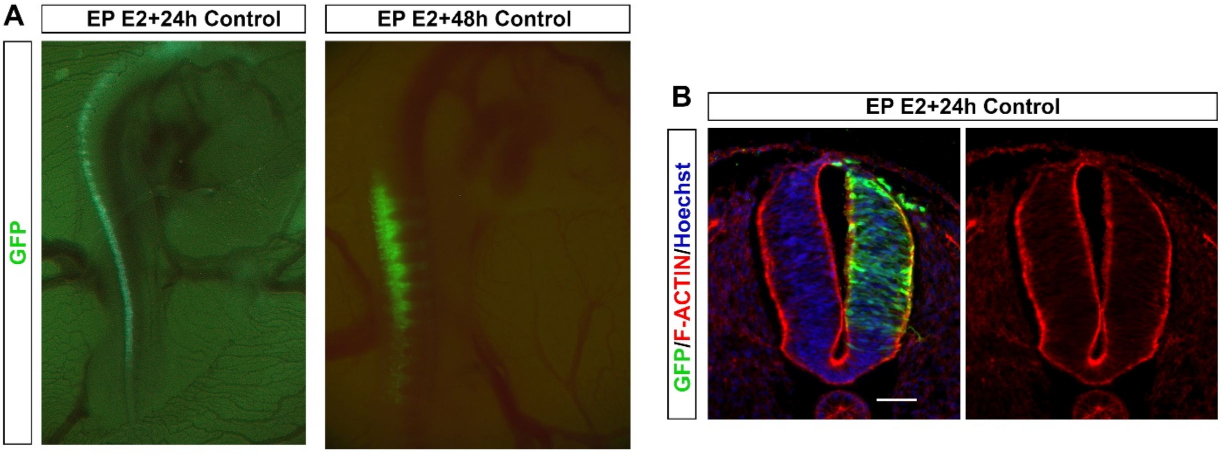


**Supplementary Figure 6:**

**A -** Dorsal view of whole-mount embryo one day and two days after the electroporation of the control plasmid (pCAGGS, expressing only GFP) using a fluorescent binocular microscope, highlighting GFP+ transfected cells. **B -** Immunofluorescence staining with the anti-GFP antibody and F-ACTIN staining on trunk transverse section of chicken embryo one day after electroporation of the control plasmid. Blue represents Hoechst nuclear staining. Scale bar: 50 µm

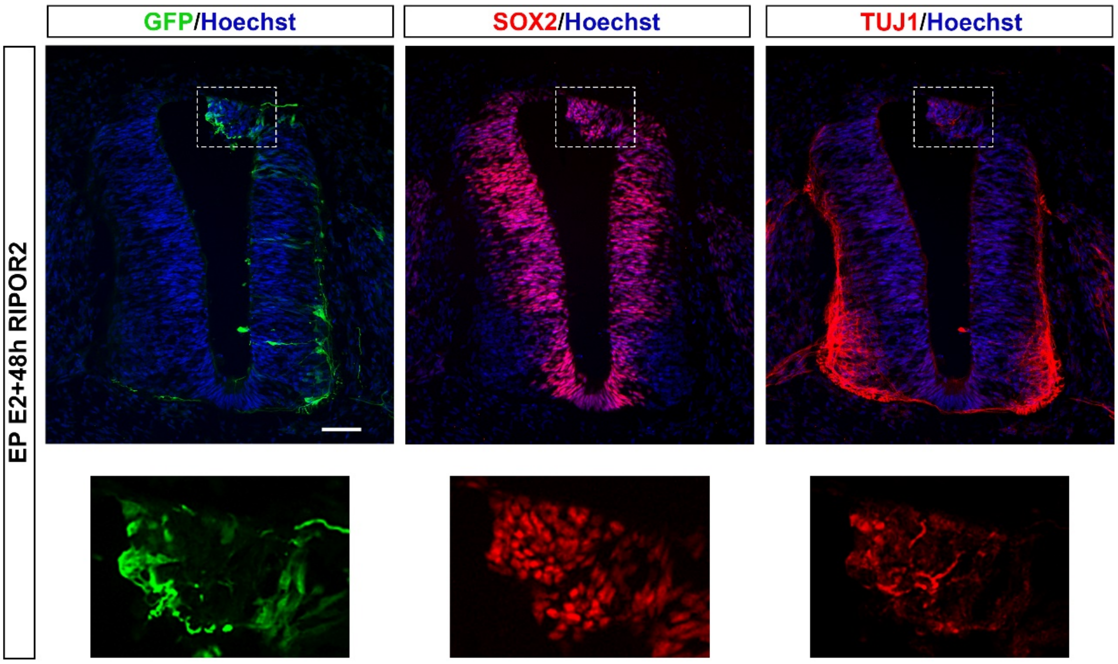


**Supplementary Figure 7:**

Immunofluorescence staining with the anti-GFP, anti-SOX2, and anti-TUJ1 antibodies on trunk transverse sections of chicken embryo two days after electroporation of RIPOR2, highlighting the disorganization of the neuroepithelium as TUJ1 staining is observed at the apical part of the neural tube. Dotted boxes are magnified in the bottom panel. Blue represents Hoechst nuclear staining. Scale bar: 50 μm

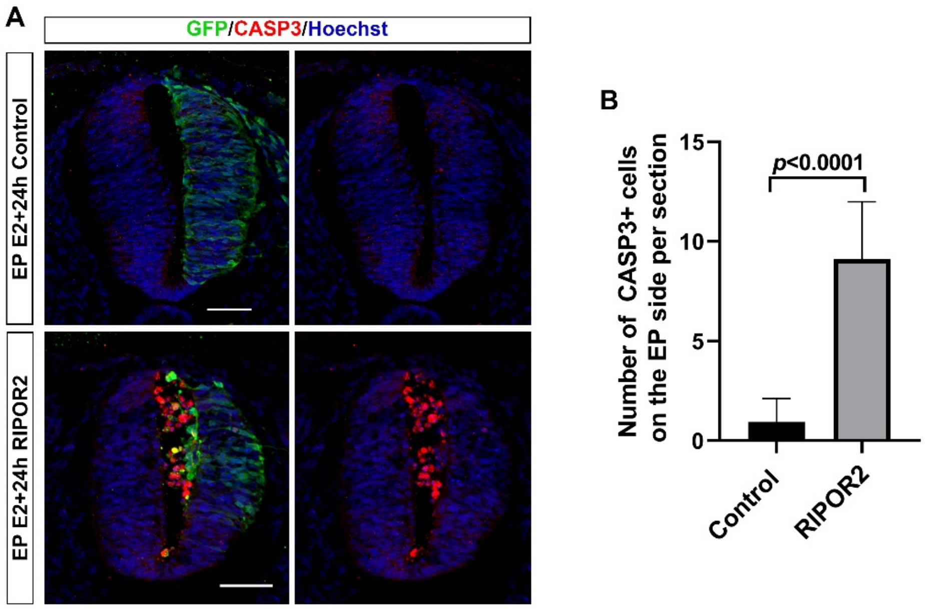


**Supplementary Figure 8:**

**A -** Immunofluorescence staining with the anti-GFP and anti-CASP3 antibodies on trunk transverse sections of chicken embryos one day after electroporation of the control vector (pCAGGS) and RIPOR2 vector. Blue represents Hoechst nuclear staining. Scale bar: 50 µm. **B -** Quantification of the number of CASP3+ cells on the electroporated side between the control (pCAGGS) and RIPOR2 conditions (n=3 animals, 18 sections, two-tailed Mann–Whitney test, error bars represent s.d.)

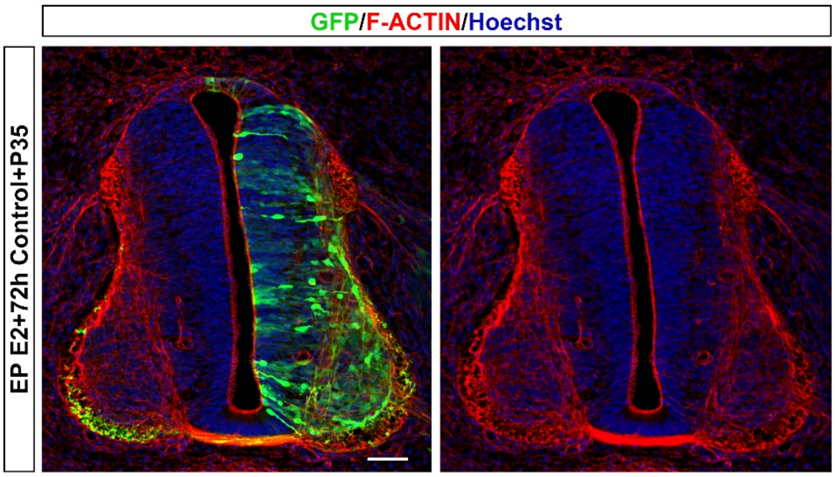


**Supplementary Figure 9:**

Immunofluorescence staining with the anti-GFP antibody and F-ACTIN staining on trunk transverse section of a chicken embryo three days after electroporation of the control vector (pCAGGS) and P35 vector. Blue represents Hoechst nuclear staining. Scale bar: 50 μm

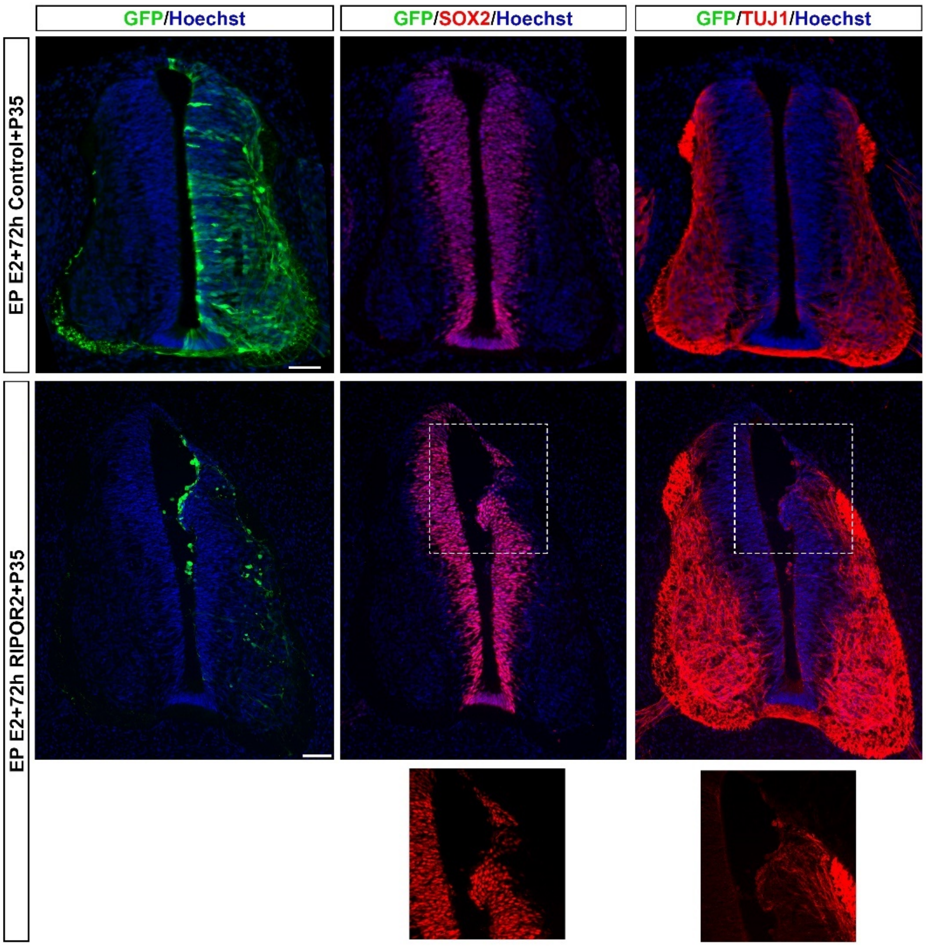


**Supplementary Figure 10:**

Immunofluorescence staining with the anti-GFP, anti-SOX2, and anti-TUJ1 antibodies on trunk transverse sections of a chicken embryo three days after electroporation of the control (P35 vector only) or RIPOR2 andP35 vectors. Reduced Sox2 and increased TUJ1 staining can be observed at the apical part of the neural tube in the RIPOR2 and P35 condition. Dotted boxes are magnified in the bottom panel.  Blue represents Hoechst staining. Scale bar: 50 µm

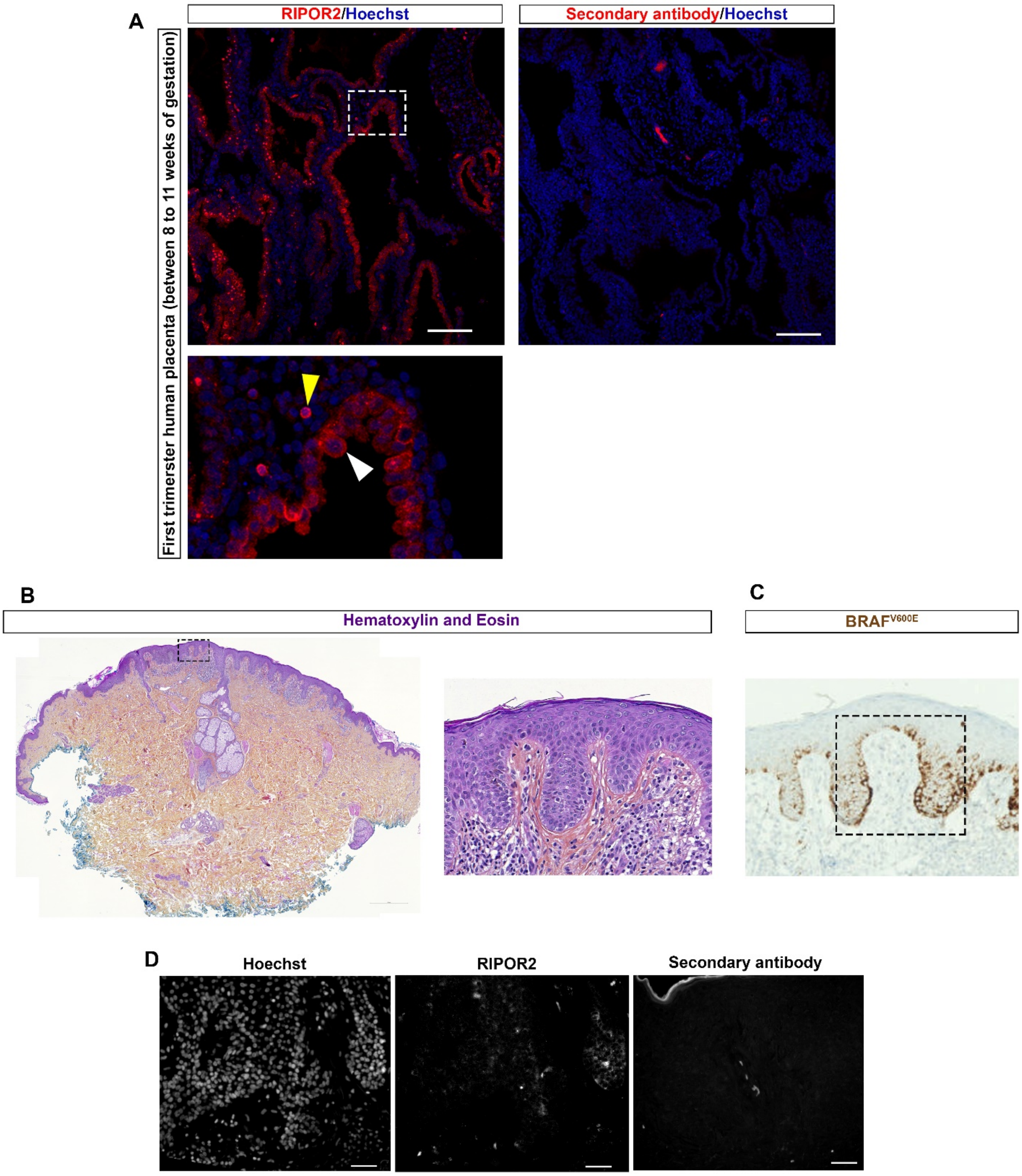


**Supplementary Figure 11: Validation of RIPOR2 antibody**

**A-** To validate the antibody, immunofluorescence staining was performed with the RIPOR2 antibody on transverse section of human placenta from the first trimester (8-11 weeks of gestation). RIPOR2 is expressed at the outer edge of the syncytiotrophoblast (white arrowhead), as previously described (Darkour et al., 1997). Note that RIPOR2 is also expressed in immune cells (yellow arrowhead). The secondary antibody alone (rabbit-647) was used as a control. Blue corresponds to Hoechst nuclear staining in all panels. **B-D-** Adjacent sections of a BRAF$^{V600E}$-benign melanocytic nevi. Dotted boxes are magnified in **D**. **B-** H&E stain shows tissue disorganization. **C-**Immunohistochemistry with an anti-BRAF$^{V600E}$ antibody stains the mutated melanocytes and highlights the malignant lesion zone. **D-** Hoechst nuclear staining and anti-RIPOR2 antibodies staining. The secondary antibody alone (rabbit-647) was used as a control. Scale bar: 100µm.

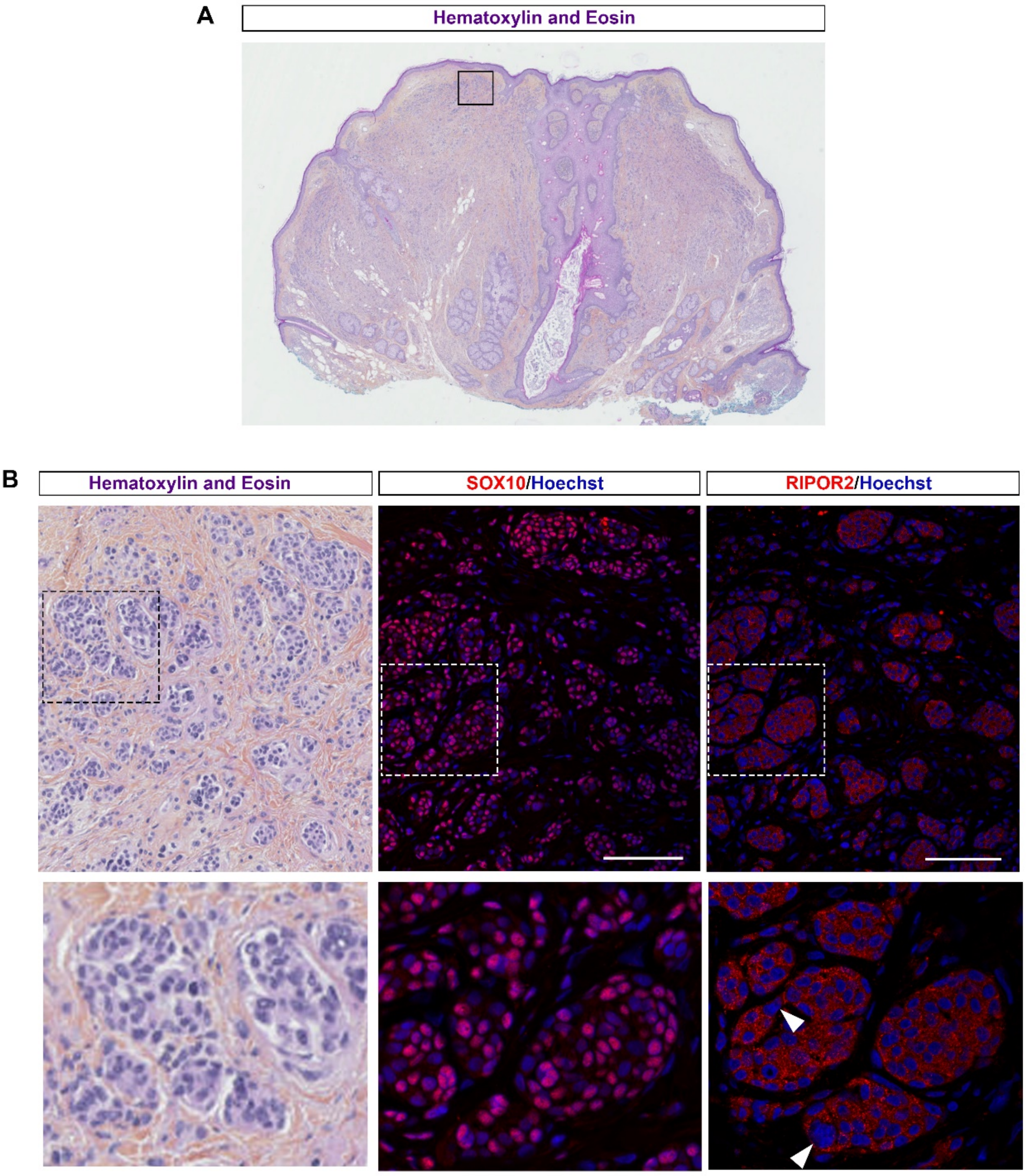


**Supplementary Figure 12:**

Adjacent sections of a benign melanocytic nevus showing multinucleated cells and neuroid maturation, $BRAF^{V600E}$ -positive. Dotted boxes are magnified in the adjacent panels. **A -** H&E stain shows tissue disorganization in the epidermis and dermis. **B** - Immunofluorescence staining with the anti-SOX10 shows SOX10 expression in melanocytes. Immunofluorescence staining with the anti-RIPOR2 antibodies demonstrates that, similar to the SOX10 melanocytes marker, RIPOR2 is expressed in the neuroid structure formed by transformed melanocytes in the dermis. White arrowheads point to RIPOR2+ multinucleated cells. Blue represents Hoechst nuclear staining in all panels. Scale bar: 100 μm

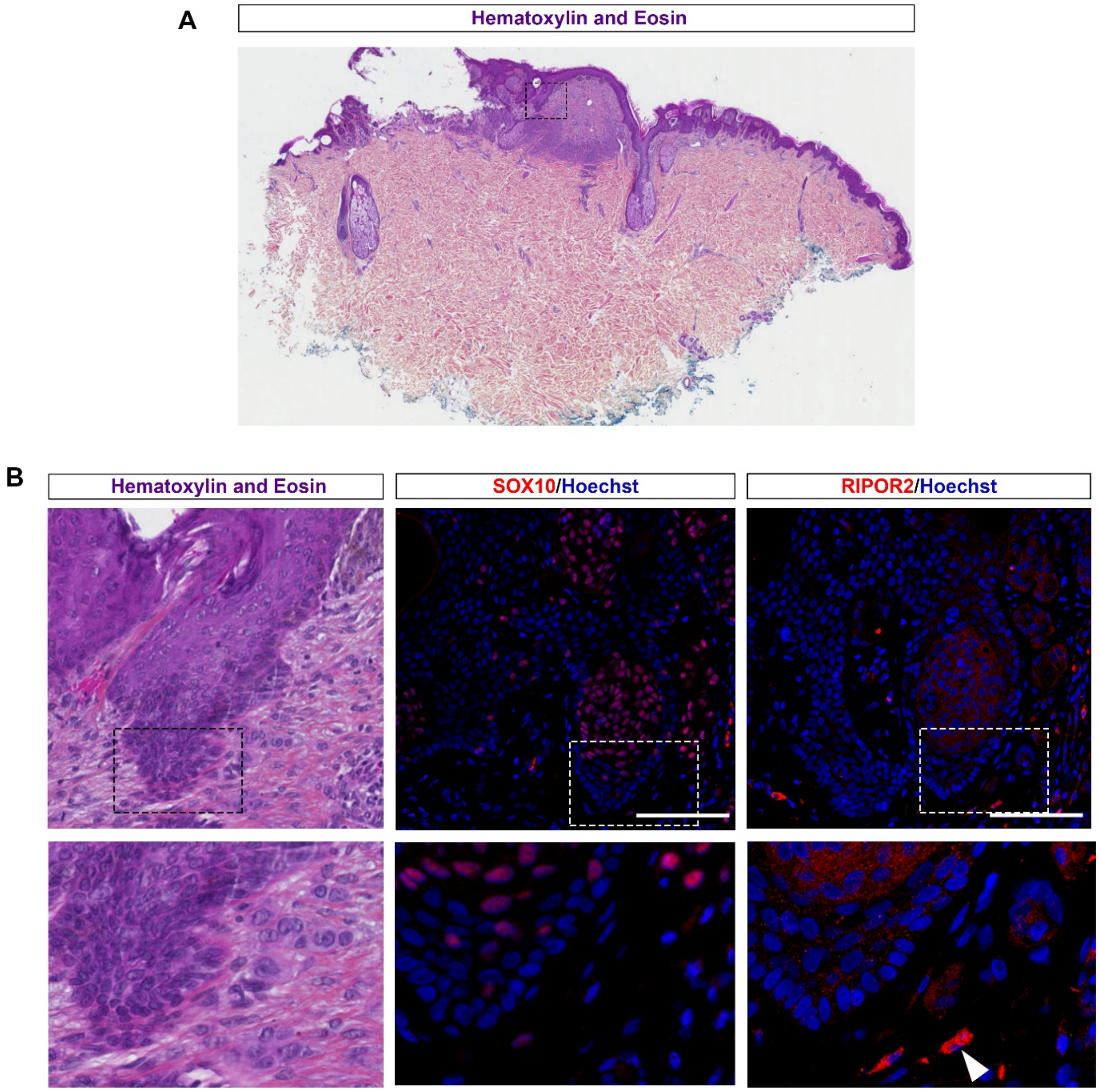


**Supplementary Figure 13:**

Adjacent sections of a dysplastic melanocytic nevus, $BRAF^{V600E}$-positive. Dotted boxes are magnified in the adjacent panels. **A -** H&E stain shows tissue disorganization in the epidermis. **B -** Immunofluorescence with the anti-SOX10 and anti- RIPOR2 antibodies demonstrates that RIPOR2 is expressed in SOX10+ structures formed by transformed melanocytes in the epidermis. White arrowheads point to an immune cell expressing RIPOR2 (higher compared to melanocytes). Blue represents Hoechst nuclear staining in all panels. Scale bar: 100 µm

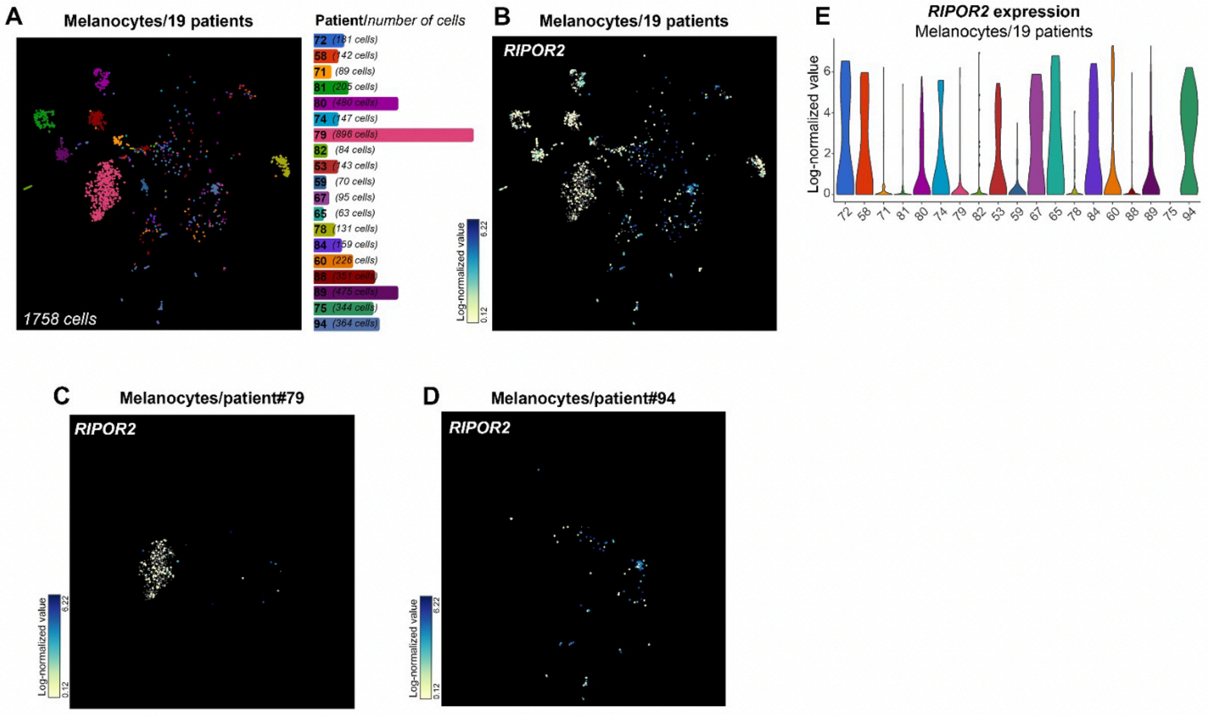


**Supplementary Figure 14: *RIPOR2* is expressed in the melanocytes of most melanoma**

Single cell RNAseq analyses using BBrowser 2 and the transcriptomic data from 19 melanoma patients from Tirosh et al., 2016 (GSE72056). **A -** t-SNE (t-distributed stochastic neighbor embedding) visualization of melanocyte cells of the 19 patients and the corresponding colour code on the right. **B-C-D** Expression of *RIPOR2* at the single cell level in this data set shows that *RIPOR2* is expressed in some of the melanocytes of most patients, with some variations (patient#79 has only a few RIPOR2 + melanocytes compared topatient#94). **E -** Violin plot of RIPOR2 expression in the melanocytes of the 19 patients. The violin plot also illustrates that *RIPOR2* is expressed in melanocytes of nearly all the patients (18/19), with strong variation.

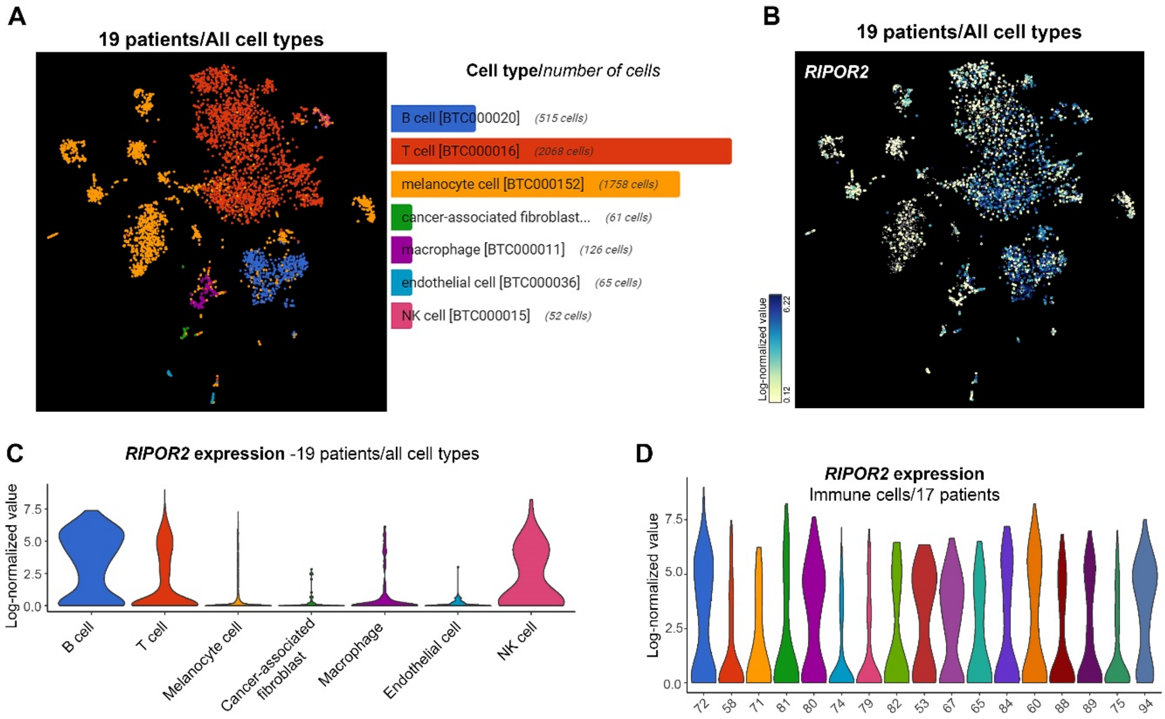


**Supplementary Figure 15:**

Single cell RNAseq analyses using BBrowser 2 and the transcriptomic data from 19 melanoma patients from Tirosh et al., 2016 (GSE72056). **A -** t-SNE visualization of all the cell types of the 19 patients and the corresponding colour code on the right. **B** – Expression of *RIPOR2* at the single cell level in this data set. **C -** Violin plot of *RIPOR2* expression in the 19 patients for different cell types shows that *RIPOR2* is highly expressed in immune cells (T cells, B cells, macrophage, and NK cells). **D -**Violin plot of *RIPOR2* expression in the immune cells of the 17 patients show that *RIPOR2* is expressed with equivalent levels of expression among them.

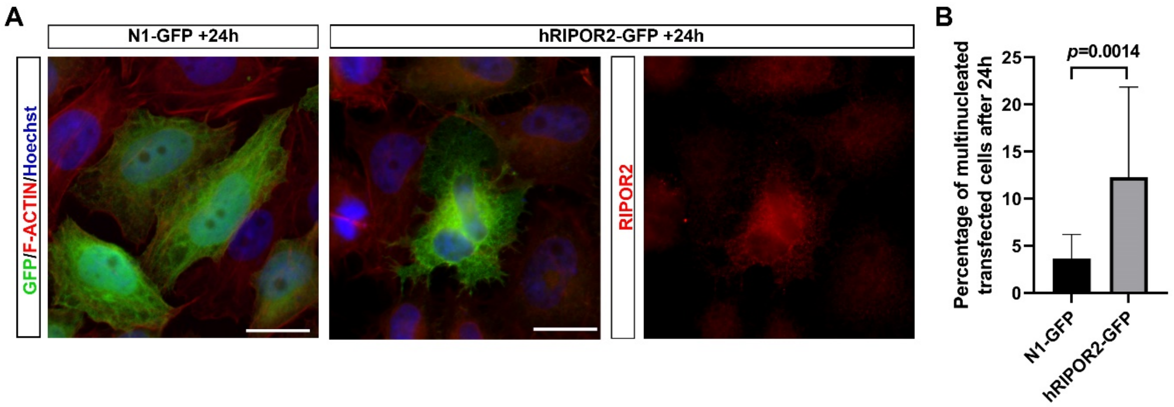


**Supplementary Figure 16:**

**A** – Immunofluorescence staining with the anti-GFP and anti-RIPOR2 antibodies and F-ACTIN staining in HeLa cell line, transfected with either a control plasmid expressing only GFP (N1-GFP) or human RIPOR2-GFP (hRIPOR2-GFP) for 24 hours. The transitory expression of h RIPOR2 increases the number of transfected (GFP+) multinucleated cells, quantify in **B -** represented as the percentage of transfected multinucleated cells (N1-GFP: 559 cells; hRIPOR2: 595 cells; 5 independent experiments, Fisher's exact test). Blue represents Hoechst nuclear staining. Scale bar: 20 μm

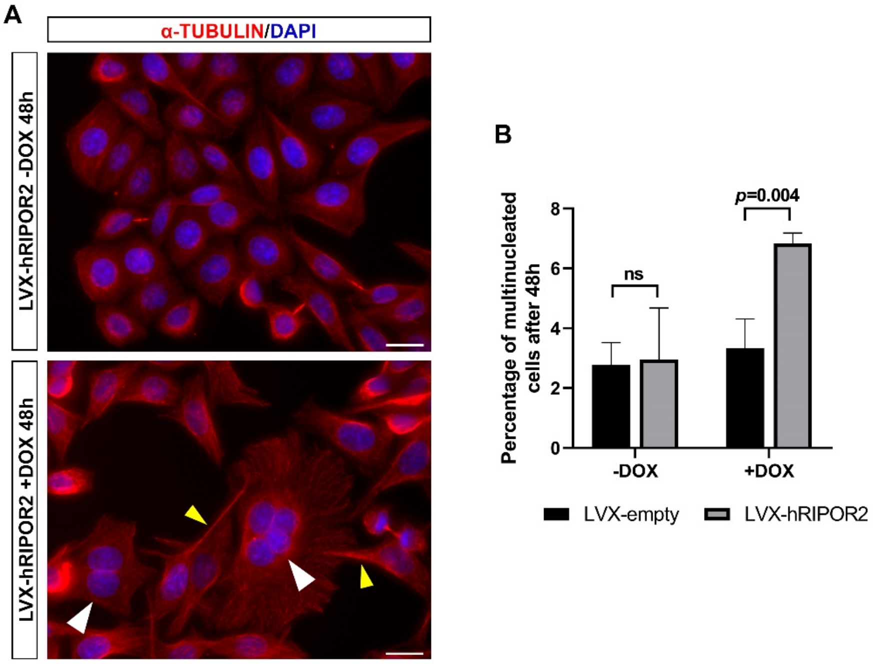


**Supplementary Figure 17:**

**A -** α-TUBULIN staining in the HeLa cell line, stably infected with doxycycline-inducible plasmid (LVX-TetOne-Puro) expressing either only GFP (LVX-GFP) or human RIPOR2 (LVX-hRIPOR2). The two cell lines were incubated with doxycycline (+DOX) or in the absence of doxycycline (-DOX, control) for 48 hours. The stable expression of hRIPOR2 induces cellular protrusions (yellow arrowheads) and increases the number of multinucleated cells (white arrowheads), quantified in **B -** represented as the percentage of multinucleated cells (LVX-GFP -DOX: 711 cells, LVX-GFP +DOX: 659 cells, LVX-hRIPOR2 -DOX: 635 cells, LVX- hRIPOR2 +DOX: 643 cells; 3 independent experiments, Fisher's exact test). Blue represents Hoechst nuclear staining. Scale bar: 20 µm.
